# Mucosal vaccine immunity induced by a new auxotrophic *Pseudomonas aeruginosa* strain is linked to Th17 and IgA responses

**DOI:** 10.1101/2025.10.03.679932

**Authors:** Y Sereme, B Villeret, S Caboche, S Ikeh, D Beury, F Maurier, C Desterke, M Born-Bony, Z Xing, R Voulhoux, Sallenave J-M

## Abstract

*Pseudomonas aeruginosa* (*P.a*) is a Gram-negative opportunistic pathogen that poses a major global health threat, particularly in immunocompromised individuals, patients with cystic fibrosis, and those with burn injuries or ventilator-associated pneumonia. Despite intense efforts, no licensed vaccine is currently available for human use. In this context, live attenuated vaccines (LAVs) represent a promising but underexplored approach, offering the potential to elicit robust, long-lasting, and multifaceted immune responses including that of inducing trained immunity. Here, we sub-cultured Δ*LasB* PAO1 (a *P.a* strain that we have shown previously shown to have reduced virulence) in artificial sputum medium (ASM), a culture medium mimicking CF sputum in which bacteria often show auxotrophy. We showed that such a strain (designed here ‘V’ for vaccine) was auxotrophic, less virulent, and had characteristics of ‘CF-like strains’. Crucially, V was able to induce both local (IgA) and systemic humoral responses as well as memory Th17 immune responses, and could, when administered intra-tracheally (but not intra-muscularly), fully protected mice against a lethal PAO1 infection. Overall, the present study demonstrates that our vaccine formulation, in addition to providing an advantageous auxotrophic phenotype adapted to the CF setting, was efficient, when given mucosally, in preferentially inducing secretory IgA and Th17 pathway at mucosal surfaces, a critical barrier that neutralizes pathogens before tissue invasion.

## INTRODUCTION

*Pseudomonas aeruginosa* (*P.a*) is a Gram-negative opportunistic pathogen that poses a major global health threat, particularly in immunocompromised individuals, patients with cystic fibrosis, and those with burn injuries or ventilator-associated pneumonia. It is a leading cause of hospital-acquired infections, contributing to significant morbidity and mortality worldwide. The World Health Organization has classified *P.a* as a critical priority pathogen due to its remarkable intrinsic resistance and its ability to rapidly acquire new antibiotic resistance determinants. Despite the fact that the use of CFTR correctors and potentiators has a very important clinical impact in CF (1–2), the rise of multidrug-resistant (MDR) and extensively drug-resistant (XDR) strains is still severely restricting therapeutic options and underscores the urgent need for effective preventive strategies. Despite this, no licensed vaccine is currently available for human use. Although various platforms—including subunit, conjugate, and outer membrane protein-based vaccines—have been explored, most candidates have failed to provide broad, durable protection in clinical trials. In this context, live attenuated vaccines (LAVs) represent a promising but underexplored approach, offering the potential to elicit robust, long-lasting, and multifaceted immune responses including that of inducing trained immunity (3–7).These include strong humoral and cellular (CD4⁺/CD8⁺ T-cell) immunity—that closely parallels natural exposure (8). Additionally, certain live vaccines (e.g., oral polio, rotavirus) administered mucosally stimulate localized IgA and mucosal immunity, which can reduce transmission—a benefit rarely seen with inactivated or subunit formulations (9). Their replicative nature enables dose sparing, reducing antigen requirements and facilitating cost-effective mass production and distribution (10).

In the context of vaccination against *P.a*, it was shown that by engineering auxotrophic bacterial strains that lack the ability to synthesize key nutrients—such as aromatic amino acids (*ΔaroA*) or D-glutamate—such live attenuated vaccines can be rendered replication-limited outside of controlled environments (11–13). Such auxotrophic mutants retain immunogenic components required to stimulate strong mucosal and systemic immunity, including Th17 and IL-17 responses, while being incapable of causing disease (12–13). In that context, our rationale was to elaborate, for the first time to our knowledge, a specific auxotrophic *P.a* strain phenotypically as close as possible to clinical strains present in sputa secretions from patients with CF (pwCF), with reduced virulence while demonstrating adequate immunogenicity.

Here, we sub-cultured ***Δ****LasB* PAO1, a *P.a* strain that we have shown to have reduced virulence (14–15) in artificial sputum medium (ASM), a culture medium mimicking CF sputum in which bacteria often show auxotrophy (16–20). Indeed, we show that such a strain (designed here ‘V’ for vaccine) was auxotrophic, less virulent, and had characteristics of ‘CF-like strains’. Crucially, V was able to induce both local (IgA) and systemic humoral responses as well as memory Th17 immune responses, and could, when administered intra-tracheally or intra- nasally, fully protect mice against a lethal PAO1 infection.

## METHODS

### Ethical statement

Animal experiments were carried out at the animal facility of the Paris-Cité Faculty of Medicine, Bichat site in Paris, in accordance with current regulations (Directive 2010/63/EU of 22 September 2010 and Decree no. 2013-118 of 1 February 2013 on the protection of animals used for scientific purposes). The protocol was approved by an animal experimentation ethics committee approved by the French Ministry of Higher Education and Research (APAFIS 27834).

### Bacterial strains and culture

PAO1-WT and clinical strains are described in Table 1 and reference 21). The vaccine strain PAO1-*ΔlasB* (‘V strain’) was generated starting as follows :

**Table 1.**
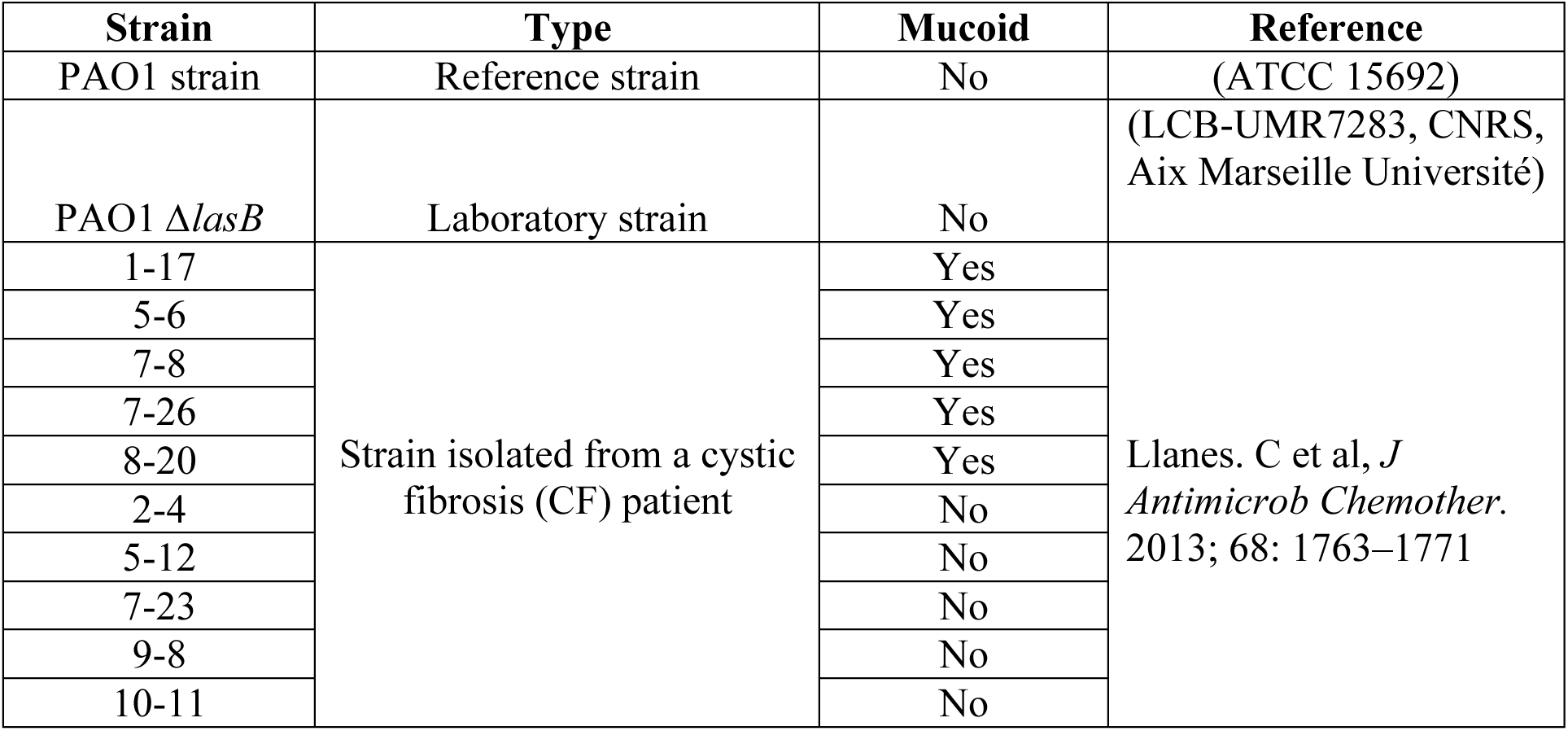
Resource bacterial strains.

PAO1Δ*lasB* was constructed from PAO1 wild type strain (22), using overlapping PCRs with specific primers. Homologous recombinations were carried out between the 5’ (∼500-bp) and 3’ (∼500-bp) regions flanking *lasB* using the pKNG101Δ*lasB* plasmid and according to the protocol of (23). To construct the pKNG101Δ*lasB*, two DNA fragments corresponding to the ∼500-bp upstream and ∼500-bp downstream regions of *lasB* gene (PA3724) were amplified from PAO1 chromosomal DNA by PCR and ligated by overlapping PCR and cloned into linearized pKNG101 by the SLIC method. The resulting constructs were transformed into *E. coli* CC118λpir and introduced into *P. aeruginosa* PAO1 by conjugation. The strains in which the chromosomal integration event occurred were selected on *Pseudomonas* isolation agar gentamycin (Gm) plates. Excision of the plasmid, resulting in the deletion of the chromosomal target gene was performed after selection on Luria-Bertani (LB) plates containing 6% sucrose. Clones that became sucrose resistant and Gm sensitive were confirmed to be deleted for the *lasB* gene by PCR analysis.

The PAO1-Δ*lasB* ‘V’ strain was then obtained by sequential growth over 15 days in artificial sputum medium (ASM), by changing the ASM medium every four days.

Before experiments, this ‘V’ strain and other *P.a* strains (PAO1-WT, PAO1- *ΔlasB*, PAO1- *ΔlasB*-HI (heat-inactivated), mucoïd and non mucoïd clinical strains, all cultured in LB medium) were grown overnight at 37 °C on LB agar plates. Single colonies were then inoculated into 10 mL of LB broth and incubated overnight at 37 °C with shaking (180 rpm) to prepare a pre-culture. The optical density (OD) at 600 nm was measured, and the bacteria were then diluted in fresh LB to reach an OD of 0.08. This culture was incubated at 37 °C with shaking for 2–3 hours until it reached exponential growth, corresponding to an OD between 0.4 and 0.8. The OD was measured again before further use.

### Bacterial auxotrophy test

The various PAO1 strains were cultured until they reached the exponential growth phase (DO between 0.4 and 0.8), and the optical density (DO) was measured at 600 nm. The bacteria were then resuspended in 10 mL of laboratory-made M9 minimal medium (Gibco by Life Technologies) to obtain an OD of 0.05. Incubation was carried out at 37°C with agitation (210 rpm). DO was measured every two hours for bacteria grown in M9 medium and the vaccine ‘V’ strain was also, when needed, grown in M9 medium supplemented with aromatic amino acids (40 µg/mL tryptophan, tyrosine and phenylalanine), and DO was measured as above.

### Pyocyanin assay

One milliliter of overnight bacterial culture from PAO1 strains was centrifuged to pellet cells, and the O.D of the supernatant was measured at 625 nm for the assessment of pyocyanin production.

### Bacterial motility assays

PAO1 strains were grown to exponential phase (OD₆₀₀ = 0.4–0.8). Swimming and swarming were assessed by spotting 10 µL of culture at OD₆₀₀ = 0.1 on 0.3 % agar plates. Plates were incubated at 37 °C for 8 h (swimming) or 24 h (swarming), and motility zones were measured. Contraction motility was evaluated by spotting 10 µL at OD₆₀₀ = 0.1 on 1 % agar overlying plastic. After 24 h, the agar was removed and bacterial spread on the plastic was measured.

### Cell line culture and maintenance

DC2.4 dendritic cells (24) were cultured in IMDM supplemented with L-glutamine, HEPES, 10 % FBS, and penicillin/streptomycin. Cells were seeded at 2 × 10⁵ cells/mL in 100 mm dishes and passaged weekly when suspensions reached 2 × 10⁶ cells/mL. Adherent cells were detached using PBS with 10 mM EDTA. MUTU (DC1, 24), MPI alveolar macrophages (25) and DJS-2 epithelial cells (26) were cultured similarly, using IMDM (MUTU), RPMI (MPI with 30 µg/mL mGM-CSF), or DMEM media (DJS-2). Adherent cells were recovered by EDTA or trypsin treatment, counted, and replated at 2–2.1 × 10⁵ cells/mL.

### In vitro PAO1 adhesion assay

Cells (3 × 10⁵ per well) were seeded in 16-well plates and incubated overnight at 37 °C in appropriate medium with 10 % FBS and antibiotics. PAO1 bacterial strains were prepared in media (see section above) with 10 % FBS and used to infect cells at a multiplicity of infection (MOI) = 1 during 1h. After washing thrice with PBS, cells were lysed with 500 µL PBS containing 0.02 % Triton on ice for 10 min. Released bacteria were plated on LB agar and incubated overnight at 37 °C for CFU enumeration.

### In vitro viability assay

Cells were plated in 96-well plates at 5 × 10⁴ per well and incubated overnight. Cells were infected (MOI = 1) and incubated for 3 h (MPI) or 24 h (other cell lines, see above). Ten microliters of MTT reagent were added, followed by a 4-hour incubation. Media were replaced with 50 µL solubilization solution for 30 min at 37 °C. Plates were read at 540 nm on a TECAN reader.

### In vivo experiments

#### a) Animals

Seven to ten-week old male C57BL/6 mice were purchased from Janvier Labs. Animals were kept in a specific pathogen-free facility under 12-h light/dark cycles, with free access to food and water. Procedures were approved by our ethical committee and by the French Ministry of Education and Research (APAFIS 27834).

#### b) In vivo Immune responses analysis

C57/Bl6 mice were immunized with three doses of various PAO1 strains (10^6 cfu of vaccine ‘V’ PAO1 strain, PAO1-WT, PAO1- *ΔlasB*, PAO1- *ΔlasB*-HI) or PBS intra-muscularly (i.m) or via the oropharyngeal/intra-tracheal route (the two terms will be used indiscriminately throughout the manuscript) at weekly intervals, followed by collection of blood, lungs, and BAL at day 21, i.e 7 days after the last boost. Organs and body fluids were then analysed (see below) with a variety of *ex-vivo* read-outs (FACS, ELISAs…).

#### c) Mice survival after immunization and challenge

After immunization with the ‘V’ strain (or treated with PBS) as explained above, different cohorts of mice were challenged at day 21 intra-nasally (i.n) with a lethal dose of PAO1-WT (10^8 cfu) and were monitored for survival. The surviving mice (6/6 in the ‘V strain protocol’, see Fig 5) were culled at day 55 and lungs, BALs and spleens further analysed by FACS or cells were challenged *ex-vivo* with killed PAO1-WT (see below).

#### d) In vivo PAO1 clearance

24 hrs post *in-vivo* challenge with a sublethal dose (5.10^7 cfu) of PAO1-WT, lungs, BALs and spleens were collected. Organs and cells were homogeneized in PBS and homogenates were plated on TSA for CFU counts.

#### e) Measurement of anti-PAO1 specific antibodies (whole cell ELISA)

Blood and bronchoalveolar lavage fluid samples were taken 7 days after the last dose of vaccine or PBS. PAO1 WT bacteria were inactivated at 90°C for 60 minutes after overnight culture, and centrifuged at 20,000 × g for 10 min. Resuspended pellets were diluted to an DO_600_ of 0.2, and 100 µl of the different bacterial suspensions were transferred to 96-well ELISA plates, which were incubated overnight at 37°C. After blocking with 1% bovine serum albumin (BSA) and washing, 100 µL of mouse samples (serum and BAL, diluted 1/2) were added to each well and incubated for 2 h at 37°C. Next, 100 µL of a 1:10000 dilution of secondary anti-mouse antibodies (IgG (for serum and BAL) anti-H&L horse (for BAL) IgA anti-H&L goat, Abcam, Paris, France) coupled to horseradish peroxidase were added. 50 µl of substrate was then used for 30 minutes to reveal the reaction (TMB liquid substrate, TONBO Bioscience, San Diego, CA, USA) and the reaction was stopped with 50 µl of 0.2 M sulphuric acid. The reading was then taken at 450 nm using the TECAN.

### Ex-vivo tissue analysis

#### a) FACS analysis

BAL cells were pelleted, resuspended in DMEM with 10 % FBS, counted, and adjusted to 5 × 10⁶ cells/ml. After perfusion, lung tissue was digested with collagenase/DNase, filtered, centrifuged, treated with ACK lysis buffer, washed, and adjusted to adjusted to 5 × 10⁶ cells/ml. Spleen cells were homogenized in a single-cell suspension through mechanical disruption. Briefly, the spleens wre grinded, filtered on a 40µm sterile filter and adjusted to 5 × 10⁶ cells/ml. All cells were first incubated (10 min, 4°C) with mixture of a viability dye and Fc Block antibody, washed (2,000 rpm, 5min) with PBS and then incubated (30 min, 4°C) with a cocktail of cell surface conjugated antibodies in FACS buffer (PBS, 1% BSA, 0.5Mm EDTA) with either a myeloid or lymphoid antibody panel (see Table 2 for a list of antibodies and Figs S1 and S2-S3 for FACS myeloid and lymphoid lung cells gating strategies, respectively ; NB : the gating strategies for BAL cells is identical, not shown). For intra-cellular IL-17 labelling, 10⁶ lung cells were stimulated with PMA/Ionomycin in the presence of monensin and brefeldin A for 4 h. Data were acquired the same day with a LSR Fortessa cytometer (BD Biosciences) with BD FACSDiva software and analysed with FlowJo (Tree Star, Ashland, OR, USA).

**Table 2.** List of antibodies used for flow cytometry experiments.

| Antibodies | Fluorochrome | Isotype | Clone | Provider | Cat number | Dilution |
| --- | --- | --- | --- | --- | --- | --- |
| CD45 | PE/Cyanine7 | Rat IgG2b, $\kappa$ | 30-F11 | Biolegend | 103113 | 1/200 |
| CD11b | PerCP/Cyanine 5.5 | Rat IgG2b, $\kappa$ | M1/70 | Biolegend | 101227 | 1/200 |
| CD11c | Pacific Blue | Armenian Hamster IgG | N418 | Biolegend | 117321 | 1/200 |
| Siglec F | BV711 | Rat LOU, also known as Louvain, | E50-2440 | BD Biosciences | 740764 | 1/200 |
|  |  | LOU/C, LOU/M<br>IgG2a, κ |  |  |  |  |
| I-A/I-E | APC | Rat IgG2b, κ | M5/114.<br>15.2 | Biolegend | 1076<br>13 | 1/200 |
| Ly6G | Alexa Fluor<br>700 | Rat IgG2a, κ | 1A8 | Biolegend | 1276<br>21 | 1/200 |
| Ly6C | PE | Rat IgM, κ | AL-21 | BD<br>Biosciences | 5605<br>92 | 1/400 |
| Fixable<br>Viability Dye | eFluor 506 |  |  | eBioscience | 65-0866<br>-14 | 1/1000 |
| TruStain<br>FcX™ (anti-mouse<br>CD16/32) |  | Rat IgG2a, λ | 93 | Biolegend | 1013<br>20 | 1/200 |
| Mixed extra-myeloid cell marking |  |  |  |  |  |  |
| CD45 | PE/Cyanine7 | Rat IgG2b, κ | 30-F11 | Biolegend | 1031<br>13 | 1/200 |
| CD19 | Pacific Blue | Rat IgG2a, κ | 6D5 | Biolegend | 1155<br>26 | 1/200 |
| CD3ε | APC | Armenian<br>Hamster IgG | 145-2C11 | Biolegend | 1003<br>11 | 1/100 |
| NK1.1 | Brilliant Violet<br>650 | Mouse IgG2a, κ | PK136 | Biolegend | 1087<br>35 | 1/100 |
| CD335<br>(NKp46) | FITC | Rat IgG2a, κ | 29A1.4 | Biolegend | 1376<br>06 | 1/200 |
| CD4 | Alexa Fluor<br>700 | Rat / IgG2a,<br>kappa | RM4-5 | eBiosciences | 56-0042<br>-82 | 1/200 |
| CD8 | PerCP/Cyanine<br>5.5 | Rat IgG2a, κ | 53-6.7 | Biolegend | 1007<br>33 | 1/200 |
| CD69 | PE-Dazzle | Armenian<br>Hamster IgG | H1.2F3 | Biolegend | 1045<br>36 | 1/200 |
| CD44 | PE | Rat IgG2b, κ | IM7 | Biolegend | 1030<br>24 | 1/200 |
| CD138 | Brilliant Violet<br>605 | Rat IgG2a, κ | 281-2 | Biolegend | 1425<br>15 | 1/200 |
| CD62L | APC/Cyanine7 | Rat IgG2a, κ | MEL-14 | Biolegend | 1044<br>28 | 1/200 |
| Fixable Viability Dye | eFluor 506 |  |  | eBioscience | 65-0866-14 | 1/1000 |
| TruStain FcX™ (anti-mouse CD16/32) |  | Rat IgG2a, λ | 93 | Biolegend | 101320 | 1/200 |
| Mixed marking Mixed extra-cell lymphoid |  |  |  |  |  |  |
| CD45 | Pacific Blue | Rat IgG2b, κ | S18009F | Biolegend | 157212 | 1/200 |
| CD11b | FITC | Rat IgG2b, κ | M1/70 | Biolegend | 101205 | 1/200 |
| IA-IE | FITC | Rat IgG2b, κ | M5/114.15.2 | Biolegend | 107605 | 1/200 |
| CD3ε | APC | Armenian Hamster IgG | 145-2C11 | Biolegend | 100311 | 1/100 |
| CD4 | Alexa Fluor 700 | Rat / IgG2a, kappa | RM4-5 | eBiosciences | 56-0042-82 | 1/200 |
| CD8 | PerCP/Cyanine 5.5 | Rat IgG2a, κ | 53-6.7 | Biolegend | 100733 | 1/200 |
| CD25 | Brilliant Violet 650 | Rat IgG1, λ | PC61 | Biolegend | 102037 | 1/200 |
| Fixable Viability Dye | eFluor 506 |  |  | eBioscience | 65-0866-14 | 1/1000 |
| TruStain FcX™ (anti-mouse CD16/32) |  | Rat IgG2a, λ | 93 | Biolegend | 101320 | 1/200 |
| Mixed extracellular marking - Stimulation with PMA/Ionomycin |  |  |  |  |  |  |
| FOXP3 | PE | Mouse IgG1, κ | 150D | Biolegend | 320008 | 1/40 |
| IL-17A | Brilliant Violet 711 | Rat IgG1, κ | TC11-18H10.1 | Biolegend | 506941 | 1/40 |
| RORgt | PE-CF594 | Mouse IgG2a, κ | Q31-378 | BD Biosciences | 562684 | 1/40 |

### b) Ex-vivo antigenic recall with killed PAO1 WT and cytokine measurements

Lung (10⁶), spleen (10⁶), and BAL (2 × 10⁴) cells were plated and stimulated with gentamicin-killed PAO1 for 48 h in RPMI with 10 % FBS in a total volume of 0.2 ml. After centrifugation at 500g, supernatants were collected and analyzed for IFN-γ, IFN-α, IL-6, IL-4, and IL-17 cytokine levels by sandwich ELISA (R&D Systems).

#### c) Complement-dependent bactericidal activity assay

Bacterial strains were grown to OD₆₀₀ ≈ 0.6–0.7, diluted to 10⁴ CFU, and incubated with 10 µL of heat-inactivated (45 min at 56°C) mouse serum or BAL. Then, 10 μl of 3- to 4-week-old rabbit complement (Pel-Freez® Biologicals, AR, Rogers, USA) and 10 μl of PBS were added in a 96-well plate, then incubated at 37°C during an hour, and plated on LB agar o/n at 37°C prior to CFU counting (27–28).

### Data analysis

Statistical analyses were performed using GraphPad Prism v10. Results are expressed as mean ± SD. Significance was determined using two-tailed Mann-Whitney tests, one-way ANOVA with multiple comparisons, or Mantel-Haenszel log-rank tests, with p < 0.05 considered for statistical significance.

## RESULTS

### Phenotypic and auxotrophic characterization of the ‘V strain’ : Δ*LasB-PAO1* cultured in ASM medium

We cultured the Δ*LasB-PAO1* strain for 15 days in ASM, with medium changes every four days. This is a culture medium mimicking CF sputum in which bacteria often show auxotrophy (29) and we refer to the Δ *LasB*-*PAO1* strain cultured in this medium as ‘V’ (for vaccine strain) for the rest of the manuscript. We showed here that the ‘V’ strain indeed showed an auxotrophic phenotype when cultured in the minimal medium M9, and only adding specific aminoacids was able to restore its growth to a level comparable to that of the WT-PAO1 (Fig 1A). Furthermore, the phenotypic shape of the ‘V’ colonies, and their mobility behaviour (twitching, swimming, swarming) was akin to the small colony variant (scv) typically found in CF secretions (Fig 1C). In addition to auxotrophy, ‘V’ showed reduced virulence compared to PAO1 WT, as assessed by its down-regulation of pyoverdin secretion (Fig 1B) and by its diminished capacity to adhere to a variety of eukaryotic cells (alveolar macrophages, epithelial cells and dendritic cells, Fig1D). In addition, it induced less cytotoxicity in eukaryotic cells *in vitro* (Fig 1E) and was less virulent *in vivo* in C57/Bl6 mice, when given through the i.n route (not shown).

**Figure 1:**
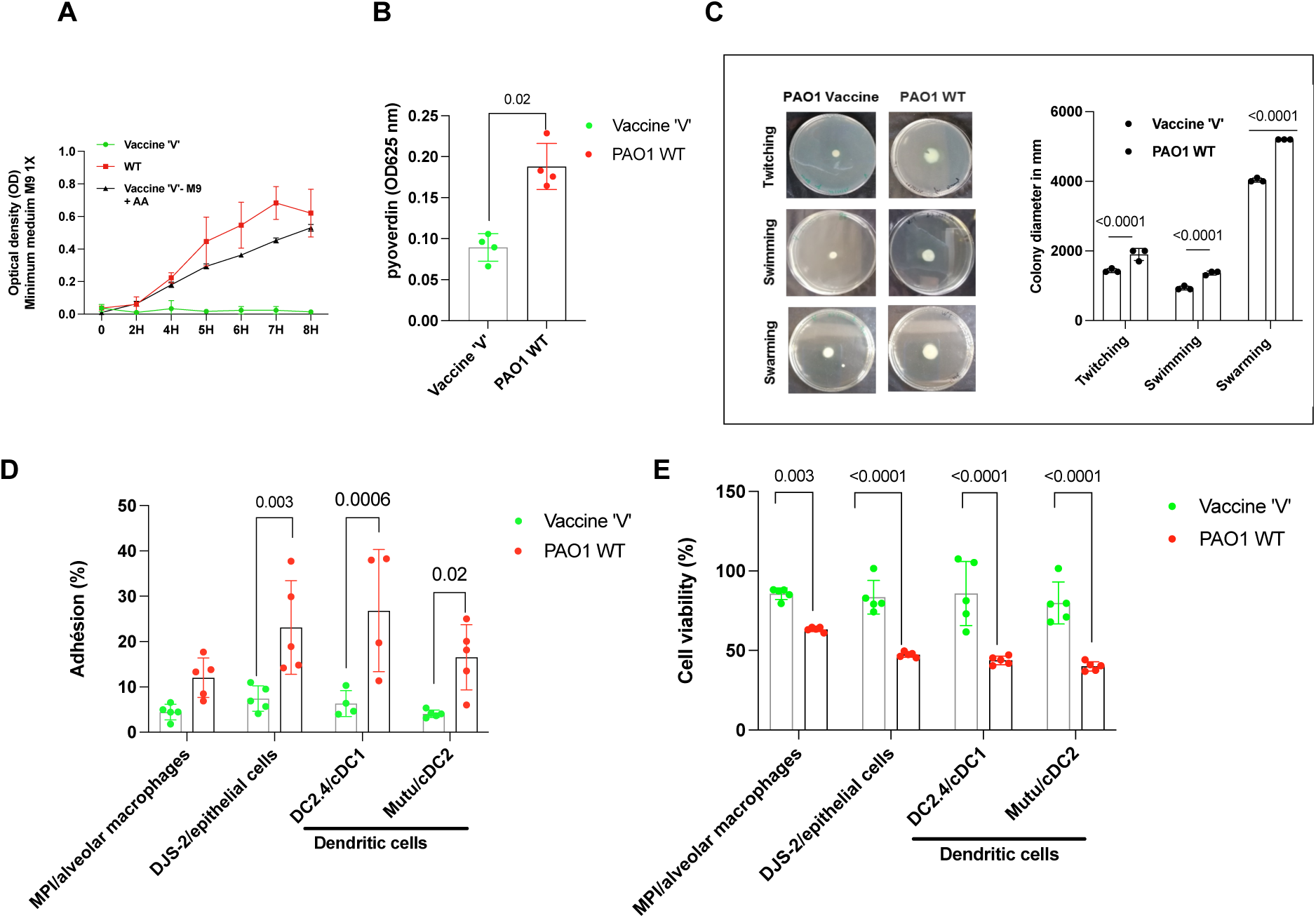
Evaluation of the virulence of the PAO1 vaccine strain ‘V’. **A)** Growth curves for *P. aeruginosa* PAO1 WT and the vaccine strain ‘V’ in M9 minimal medium and M9 supplemented with amino acids (AA). (**B**) pyoverdin production (n = 4) (**C**) measurement of motility twitching, swimming and swarming (n = 3) of the vaccine strain ‘V’, compared to the PAO1 WT strain (p < 0.05); data are presented as mean SD values. (**D**) Percentage of adhesion of PAO1 or ‘V’ to alveolar macrophages (MPI cell line, n=5), to lung club epithelial (DJS-2 cell line, n=5), and to dendritic cells DC2.4/cDC1 (n=5) and Mutu/cDC2 (n=5). Two-way ANOVA with Tukey’s multiple comparisons tests were used to determine statistical significance (p<0.05). (**E**) Cell viability of MPI (n=5), DJS-2 (n=5), DC2.4 (n=5) and Mutu/cDC2 (n=5) was measured post-infection with the bacterial strains (Two-way ANOVA with Tukey’s multiple comparisons, p<0.05).

### Assessment of the immunogenicity of the V strain

We then assessed the immunogenicity of the ‘V’ strain in C57/Bl6 mice using either a lung mucosal oro-pharyngeal route (referred indiscriminately to as ‘intra-tracheal/i.t’ for the rest of the manuscript) or an intra-muscular route (i.m) route of administration.

We showed that at day 21 (one week after the last boost), while both routes induced robust serum anti-PAO1 IgG production (Fig 2A, E), only i.t mmunization elicited high levels of specific anti-PAO1-IgG and -IgA in bronchoalveolar lavages (BALs, Fig 2 C-D), when compared to i.m vaccination (Fig 2F-G). In the latter compartment, all the tested isotypes (IgG1, IgG2a and IgG2b) were equally induced in the i.t protocol (Fig 2B).

**Figure 2:**
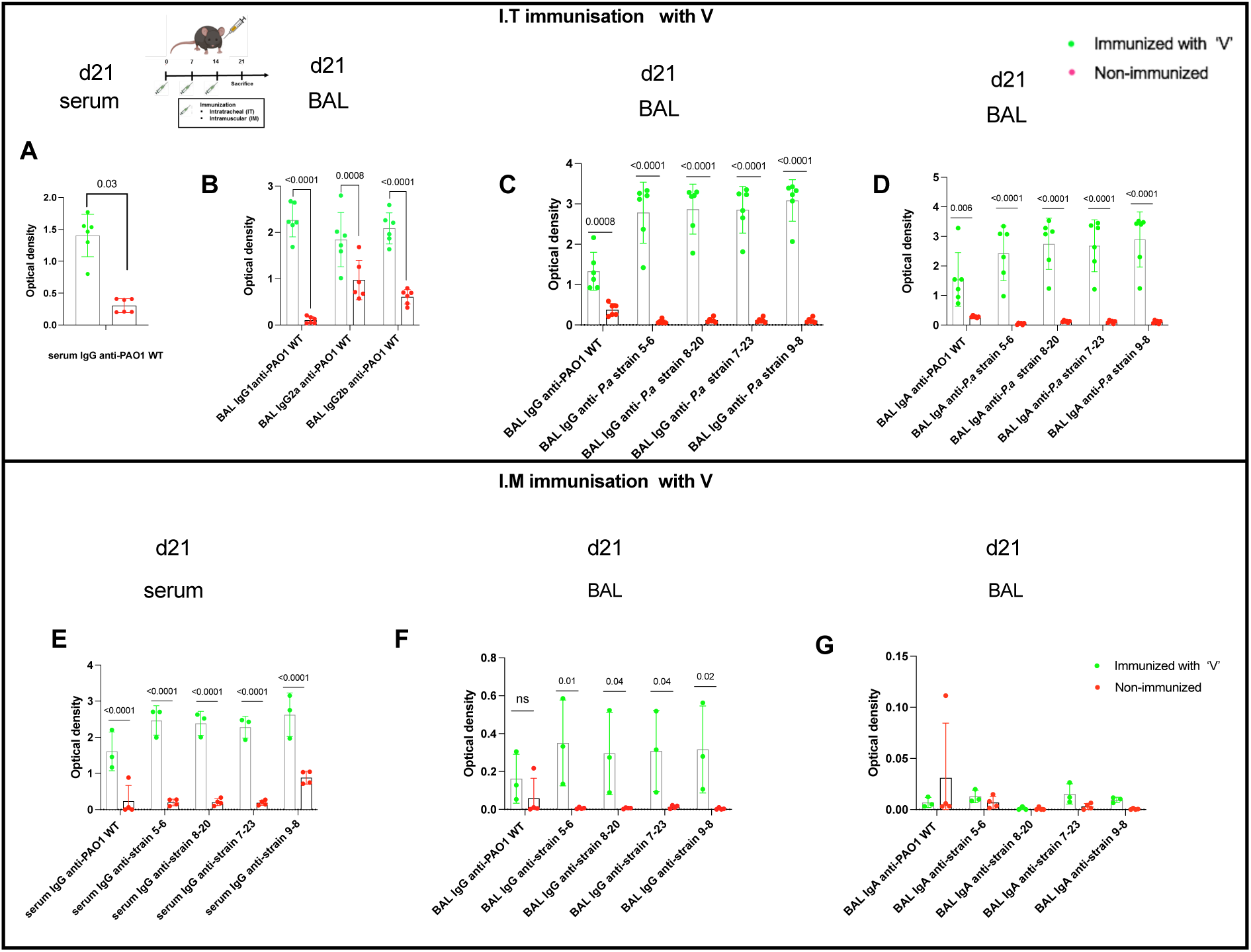
Assessment of the immunogenicity of the V strain : humoral response. Immunization protocol : At days 0, 7, 14, C57BL/6 male mice were given (IT (n=6) or IM (n=3)) either NaCl (control) or were immunized with 10^6^ CFU of the vaccine ‘V’ PAO1 strain. At day 21, 7 days after the last boost, BALs and blood were collected. (**A,E**) : Measurement of anti-PAO1, -*P.a* 5-6, 8-20, 7-23, 9-8 serum IgG levels. (**B**) Measurement of anti-PAO1 IgG1, Ig2a and IgG2b BAL levels. **(C, F)** : Measurement of anti-PAO1 BAL IgG levels. **(D,G)** : Measurement of anti-PAO1, anti-*P.a* 5-6, 8-20, 7-23, 9-8 BAL IgA levels. Statistics : Mann– Whitney non-parametric test for (**A)** and Multiple t tests were used for (B-G) and error bars represent standard deviations (SD).

Importantly, BAL IgG and IgA recognized both PAO1 WT as well as unrelated mucoid and non-mucoid clinical strains (respectively labelled 5-6; 8-20 and 7-23; 9-8, Fig 2C-D), and serum IgGs had the same cross-reactive properties (Fig 2E).

When cellular responses were assessed at the same time point (Fig 3A), mucosal immunization with ‘V’ was again more efficient at inducing immune responses in both lungs (Fig 3B) and BAL compartments (Fig 3D). Notably the numbers of neutrophils, dendritic cells (both cDC1 and cDC2), B cells, plasmocytes and CD4 T cells (both classical and tissue-memory resident/TRM) as well as those of CD8+ T cells were increased in lungs. Importantly, the proportion of IL-17+ cells among CD4+ T cells was increased in ‘V-immunized’ animals after PMA/ionomycin stimulation (Fig 3C).

**Figure 3:**
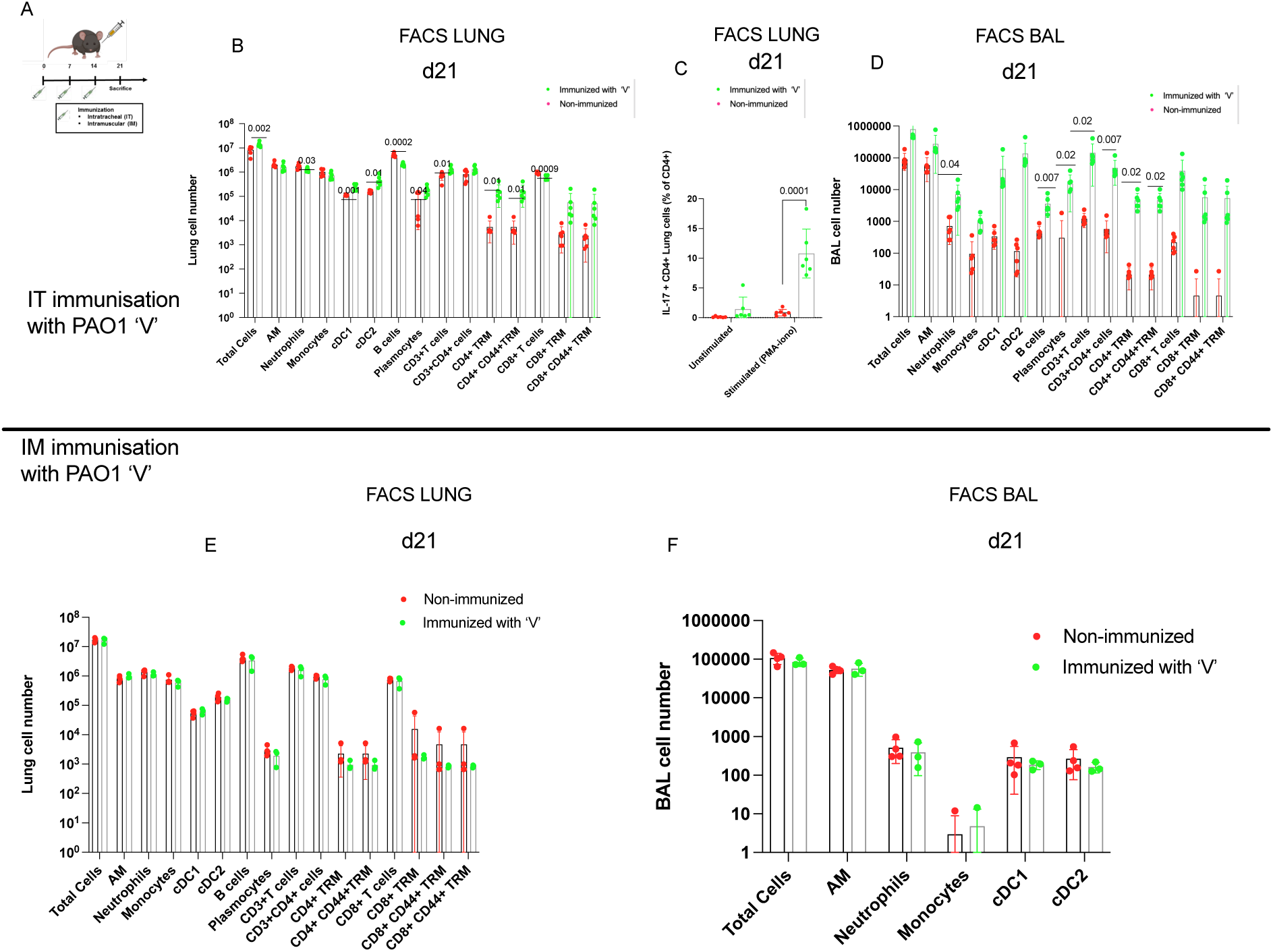
FACS assessment of the immunogenicity of the V strain; cellular response. **A)** Immunization protocol as in Fig 2. Counts of total cells, AMs, neutrophils, monocytes, dendritic cells (cDC1 and cDC2), B lymphocytes, plasma cells cells, CD3+ T cells, CD3+CD4+ T cells, CD4+CD44+TRM, CD8, CD8+TRM and CD8+CD44+TRM T cells were assessed by FACS in lungs (**B, E**) and BALs (**D, F**). (**C**): Cells were stimulated with PMA-ionomycin and % of IL-17+CD4+ cells was assessed. NB : TRMs are defined here as CD3+CD69+CD103+ .Significant intergroup differences in the different cell populations (p < 0.05). Multiple t tests were used to determine significance p<0.05. Error bars represent standard deviations (SD).

Indeed, by comparison, none of the lung and BAL cell populations were increased in i.m- immunized animals (versus non-immunized animals) (Fig 3E-F), and significantly there was a more than 2 log decrease in lung TRM between i.m and i.t immunization (Fig 3E v 3B).

We then compared the mucosal immunogenicity of the ‘V’ strain (cultured in ASM medium) to that of other PAO1 strains, i.e PAO1-WT, PAO1*-ΔLasB*, PAO1*-ΔLasB* HI (heat-inactivated), (all strains cultured in LB) and showed that the specific anti-PAO1 antibody responses in serum and BALs were enhanced when the ‘V’ strain was considered (Fig S4). When the cellular immune responses were compared, they were globally found to be equivalent, as assessed in BALs (Fig S5) and lungs (Fig S6), again demonstrating that the reduced virulence of the ‘V’ strain (Fig 1) was not affecting its immune-stimulating potential (and was even beneficial in terms of antibody responses), compared to other PAO1 strains.

We therefore focused the rest of our study on the ‘V’ strain and further showed, by performing *ex-vivo* tissue antigenic recall experiments (with killed PAO1) that mucosal i.t immunization was far superior at inducing a lung Th17 response (as measured by IL-17 levels) when compared to the i.m vaccination protocol (Fig 4A,D). Interestingly, splenic responses were equivalent in terms of IL-17 recall responses, irrespective of the route of immunization (Fig 4 C, F), and while spleen IFN-g levels were increased post i.m immunization (Fig 4F), no splenic IFN-g was detectable post i.t immunization (Fig 4C). When BAL cells antigenic recall was considered, IL-17 secretion was detectable in i.t-immunized mice (Fig 4B) but not after i.m vaccination (4E). It was likely produced by T cells, which accounted for roughly 20% of total cells in immunized mice (Fig 3D). On the other hand, BAL TNF and IL-6 were likely produced mainly by innate immune cells since these cells account for 95% of total cells in non-immunized mice and since the levels of this cytokine are roughly comparable in non-immunized and i.t immunized mice (in the latter, resident AMs and dendritic cells account for to 40% and 20% of total cells, respectively).

**Figure 4:**
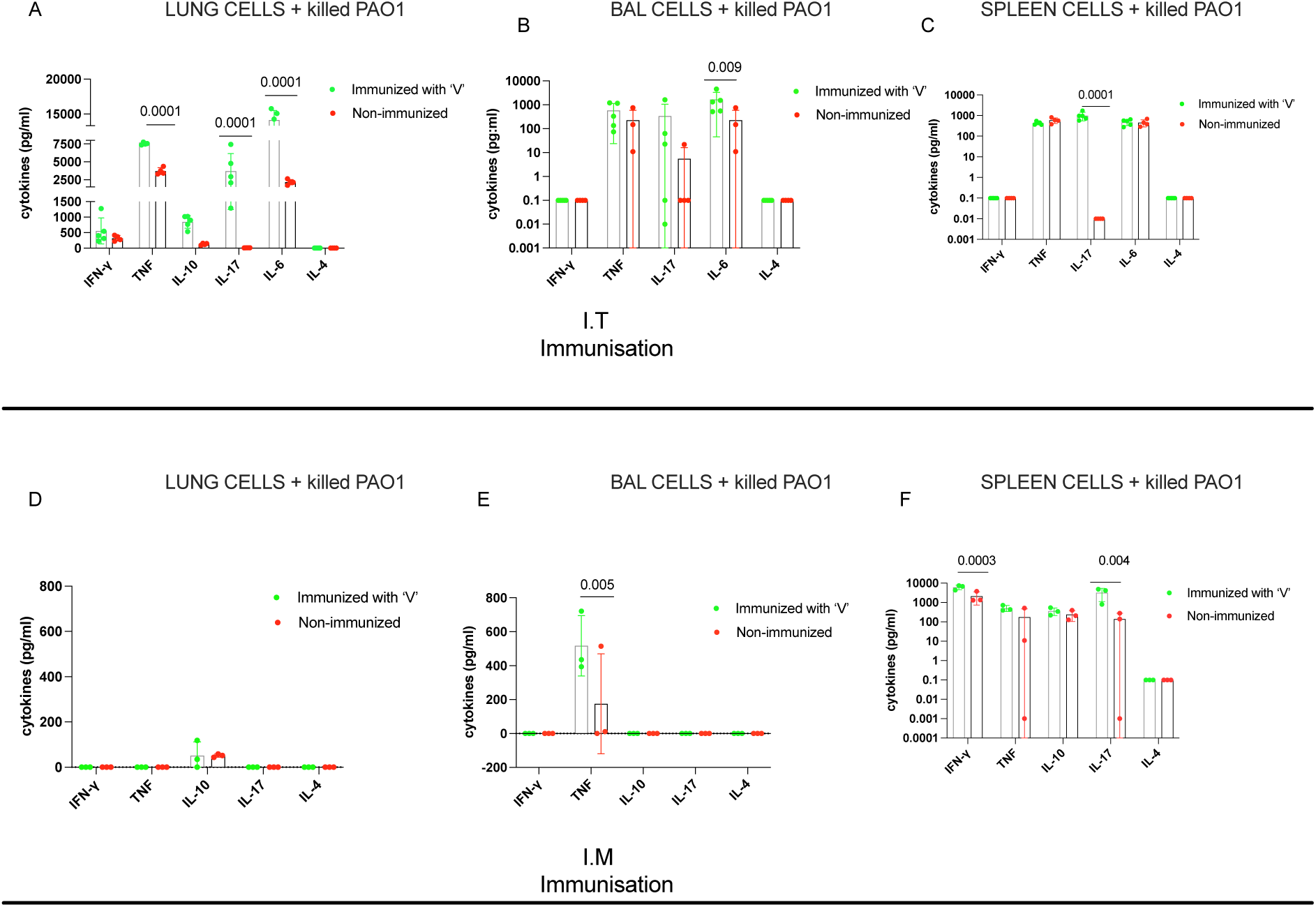
Cytokine read-out after *ex-vivo* stimulation with killed PAO1-WT. After i.t or i.m immunization (see above), lung (A,D) , spleen (C, F) and BAL (B,E) cells were harvested and cells were stimulated with gentamicin-killed PAO1 WT. IFN-g, TNF, IL-10, IL- 17; IL-6 and IL-4 cytokines levels were measured in cell supernatants 48hrs later. Two way ANOVA comparaison tests were used for statistical analysis (p<0.05). Error bars represent standard deviations (SD).

### PAO1-WT challenge experiments following immunization with the ‘V’ strain

After demonstrating above the marked differences (and few similarities) between the i.t and i.m vaccination protocols in terms of immunogenicity, we then tested how this might affect protection against a lethal *PAO1-WT* (10^8 cfu) challenge, performed intra-nasally (i.n) 7 days after the last boost (Fig 5A). We showed that i.t immunization was clearly more efficient (5B) than that of i.m at protecting mice (5C). Then, in different cohorts of mice, we showed, using a sub-lethal dose of 5.10^7 cfu of *PAO1-WT* as challenge (Fig 5D) that in i.t- immunized animals, 24hrs post-challenge, the bacterial load was very significantly diminished in the lungs of i.t-immunized (Fig 5E), but not i.m- vaccinated mice (Fig 5F).

**Figure 5:**
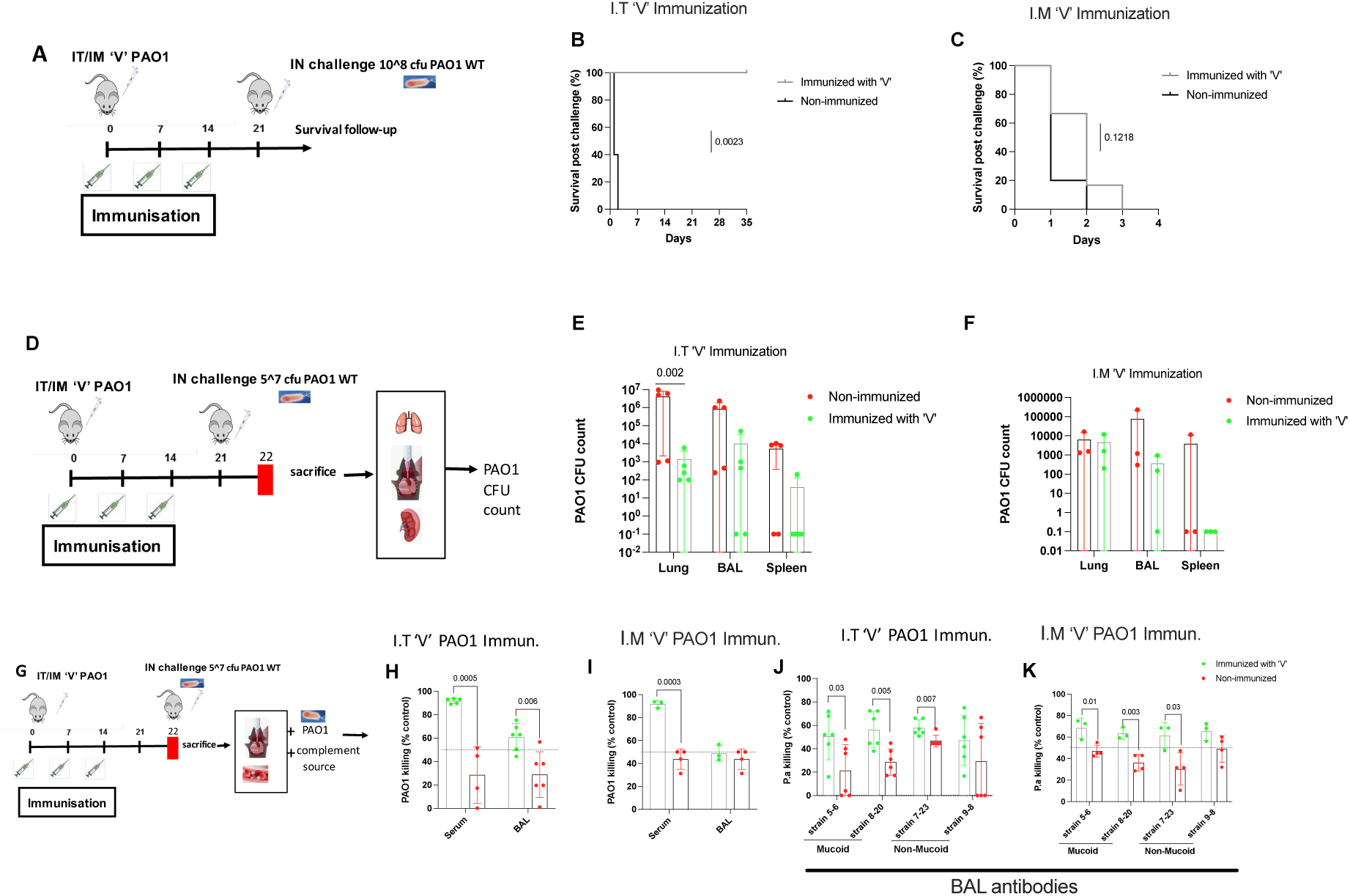
PAO1-WT challenge following either i.t or i.m immunization with the ‘V’ strain. (**A**) C57BL/6 mice were immunized i.t **(B)**, i.n or i.m **(C)** as in Figs 2 and 3. At day 21, 7 days after the last boost, mice (n=6 in each group) were challenged i.n with 10^8^ cfu of PAO1-WT. Mice were observed for survival (Mantel-Haenszel log-rank tests). (**D**) C57BL/6 mice were immunized as above (i.t (**E**) ; i.m : (**F**)), but were challenged i.n with a sub-lethal dose (5x10^7^ cfu). 24 hrs later, mice were sacrificed, and bacterial cfu counts were determined in lungs, BALs and spleens. Two way ANOVA comparaisons were used to determine statistical significance (p<0.05). Error bars represent standard deviations (SD). (**G**) : Mice were immunized as in **D)**. Sera and BALs were collected from sacrificed animals and were assessed for complement-dependent bactericidal assay. Bacterial counts were determined after incubating PAO1 WT **(H-I)** or *P.a* clinical strains **(J-K)** with either sera and BALs with an exogenous source of complement (see Materials and Methods for details). Results are expressed as % of numbers of starting PAO1 WT. Two way ANOVA comparaisons were used for to determine statistical significance (p<0.05). Error bars represent standard deviations (SD).

To assess whether antibodies may have played a role in the clearing of bacteria noted above, we then tested *ex-vivo* the complement-dependent bactericidal activity of serum and BAL antibodies against PAO1 WT and against the same *P.a* mucoid and non-mucoid clinical strains mentioned above. In keeping with data showing that both i.t and i.m immunization protocols generated strong antibody levels (Fig 2), we showed that these antibodies (from serum and BALs, Figs H-I) were able to activate the complement and kill all *P.a* strains (PAO1) and clinical strains 5-6, 8-20, 7-23, 9-8 (BAL antibodies, Figs 5 J-K), irrespective of the mode of vaccination.

### Assessment of ‘long term’ memory responses following immunization with ‘V’ and re- challenge with *PAO1*

To determine the quality of ‘long term’ memory immune responses, we assessed the status of lungs, BALs and spleens of mice at day 55 (Fig 6A), i.e 34 days after the original i.t ‘V’ immunization protocol and the ensuing i.n *PAO1* challenge (which resulted in 100% survival, see Fig 5B).

**Figure 6:**
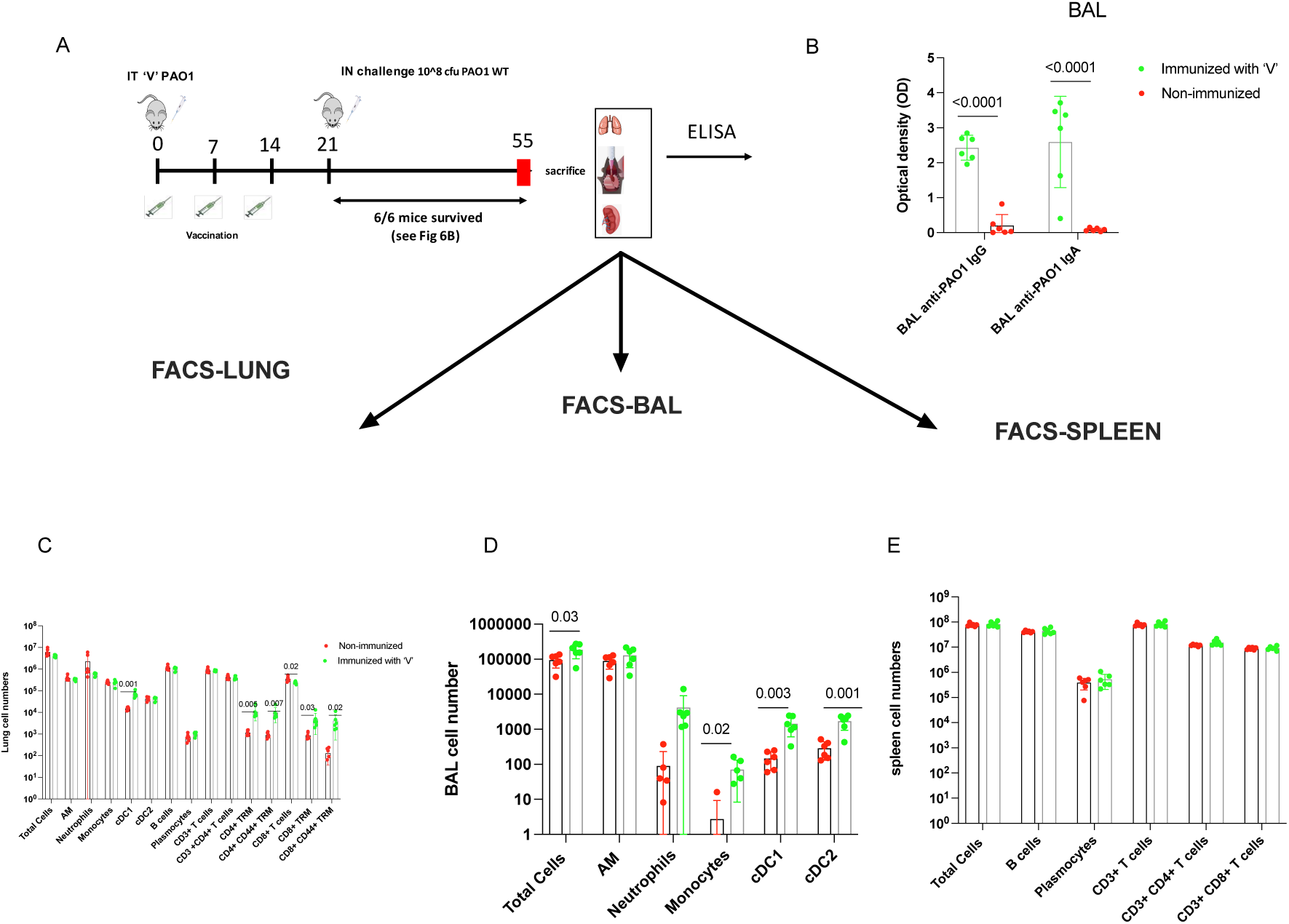
Evaluation of memory immune responses at day 55 (41 days post ‘V’ immunization and 34 days post challenge with PAO1-WT. **A)**Immunization protocol as in Fig 5. Mice which survived the challenge with PAO1 WT (6/6) were observed until day 55 (i.e 34 days post-challenge). They were then sacrificed and lungs, BALs and spleens were collected. (**B**) Anti-PAO1 WT IgG and IgA were measured in BALs. (**C-E**) Cellular immune responses were analysed by FACS as in Fig 3. Multiple t tests were used to determine statistical significance (p<0.05). Error bars represent standard deviations (SD).

Importantly, we found that the levels of specific anti-PAO1 IgG and IgA were maintained in BAL at that late time point (Fig 6B), compared to the levels observed at day 21 following ‘simple immunization’ (Fig 2C-D).

When lung immune cells were investigated, it is noticeable that although there was a slight reduction at day 55 (approximately 1 log and ½ log ) in the levels of lung CD4 and CD8 TRMs respectively, when compared to day 21 (Fig 3D), they were still very significantly increased compared to non-immunized mice (Fig 6C) In BALs, expectedly, inflammation had almost completely subsided at that time point since AMs levels returned to normal values (94.6%, Fig 6D). However, although representing minor populations in absolute numbers, monocytes, neutrophils, and DC levels were still increased in vaccinated and challenged animals at day 55 compared to untreated mice. In contrast, no differences in lymphocyte numbers were observed in the spleens between treated and control animals (Fig. 6E).

Upon antigenic recall (using the same protocol applied post-immunization on day 21; see Fig. 4), the lung parenchyma cells still secreted at that late time point (day 55) impressively high levels of IL-17 (Fig. 7C), (probably arising from the increased TRM cells described above (Fig 6C)), and these cells also showed enhanced IL-6 secretion compared to untreated animals.

**Figure 7:**
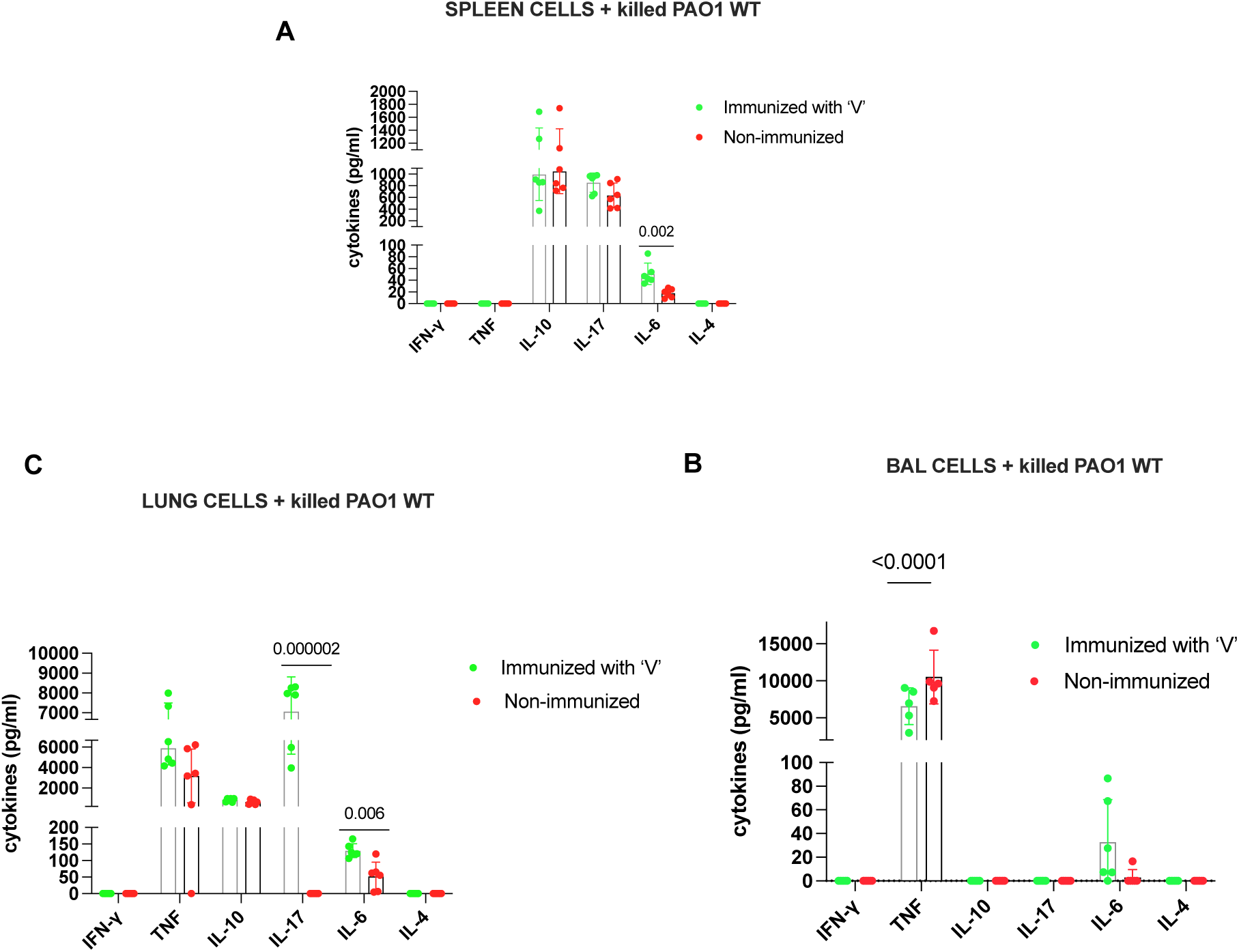
Cytokine output following long-term immunization with ‘V’ and challenge with PAO1-WT. **(A)**Immunization and challenge protocol as in Fig 6. Spleens **(A),** BALs **(B)**, and lungs **(C)** were harvested, cells were dissociated and incubated with gentamycin-killed bacteria during 48hrs. Cell supernatants were collected and IFN-g, TNF, IL-10, IL-17, IL-6 and IL-4 cytokine levels were measured. Two way ANOVA comparaisons were used to determine statistical significance (p<0.05). Error bars represent standard deviations (SD).

By contrast, splenic cells showed no difference in IL-17 levels and only a modest increase in IL-6 (Fig 7A). Interestingly, BAL cells (and most likely AMs since they were the overwhelmingly predominant cell type in BALs at day 55 (Fig. 6D)) seemed to display a memory-like phenotype and responded differently to killed PAO1 in terms of TNF (statistical significance) and IL-6 production (trend) (Fig. 7B).

## DISCUSSION

*Pseudomonas aeruginosa* (*P.a*) is an opportunistic Gram-negative pathogen responsible for severe respiratory infections, particularly in immunocompromised individuals and patients with cystic fibrosis or chronic lung disease. Its intrinsic resistance to antibiotics and ability to form biofilms make treatment challenging, highlighting the need for alternative therapeutic strategies, including prophylactic vaccines.

Here, we designed a new *P.a* vaccine strain by culturing ***Δ****LasB*-PAO1 (‘V’) in ASM, a culture medium mimicking CF sputum in which bacteria often show auxotrophy (see above). We showed that the ‘V’ strain, while indeed demonstrating an attenuated auxotrophic and reduced virulence phenotype (Fig 1), was at least as immunogenic as other PAO1 strains (WT-PAO1, ***Δ****LasB*-PAO1, ***Δ****LasB*-PAO1-HI, all cultured in classical LB medium), in terms of cellular lung and BAL recruitment (Figs S5-6), and induced higher antibody levels (Fig S4), demonstrating that auxotrophy did not impair its immune potential.

We showed that ‘V’, when given i.t, could induce both systemic and lung humoral responses (IgG and IgA, Fig 2A-D) as well as promoting lung and spleen cellular Th17 immunity (Figs 3B-C; Fig 4A-C).

Importantly, the picture was completely different when the vaccine was given intra-muscularly. Indeed, although the serum antibody response (but crucially not the BAL, Fig 2F-G) was comparable with that obtained with i.t immunization (Fig 2A,E), the lung and BAL cellular content differed drastically. Although at day 21 (7 days after the last boost), moderate inflammation was still present in BALs of i.t-immunized animals (Fig 3D), the cellular composition in the lungs of i.m- and non-immunized mice were quantitatively similar and no ‘inflammation’ was present in the BALs of either groups, with AMs by far the predominant cells (Figs 3E-F). These cell numbers differences between i.t- and i.m-immunized animals also translated functionally in antigenic recall experiments where only i.t immunization was able to generate Th17 responses (but notably neither Th1/IFN-g nor Th2/IL-4) in either lungs or BALs (Fig 4). Importantly, only that protocol significantly protected mice against a lethal challenge of PAO1 WT (Fig 5) and memory responses provided by this immunization method clearly extended to at least day 55 (Figs 6-7).

Relatedly, we and others have shown that protective immunity against *P. a* requires a coordinated response at mucosal surfaces, where Th17 cells play a pivotal role (30–34) by secreting IL-17, a cytokine crucial for neutrophil recruitment, mucosal barrier maintenance, and the induction of antimicrobial peptides (35).

In accordance, neutrophils levels were increased here in BALs after i.t (but not i.m) immunization (Fig 3D) and were still present in BALs at day 55 (Fig 6D). Interestingly, after i.t immunization, IL-6 levels were specifically enhanced even at that late time point. Whether this cytokine had a role for maintaining local Th17 differentiation (36) or as we showed before (14,37) in promoting lung repair was not assessed here. Moreover, independently of its role in protection against *P.a ,* Th17 cells have also been shown to be key players in vaccine-induced protection against Gram-negative respiratory pathogens as a whole (38–39).

Importantly, we demonstrate that our protocol efficiently fostered the generation of TRMs CD4 T cells (CD69+CD103+ cells) within the lung, which, because of their positioning, provide rapid on-site reactivity (40–44). Indeed, studies have consistently shown that mucosal delivery strategies—whether for influenza, tuberculosis, or SARS-CoV-2 antigens result in substantially higher frequencies of lung TRM compared to systemic routes. For example, live mRNA or adenoviral vectors administered via respiratory routes markedly increase CD8^+CD69^+CD103^+ T cells in lung parenchyma, while i.m immunization primarily induces circulating effector or central memory subsets (45).

Importantly, while i.n administration is widely used in both preclinical and clinical settings due to its non-invasive nature and ease of use (eg Flumist/Fluenz Tetra for *Influenza* vaccination or ChAdOx1 nCov-19 for SARS-CoV-2 Phase I trials) the i.t/.oropharyngeal route may offer distinct advantages in murine models and human settings (46–50). Specifically, i.t. delivery enables more precise targeting of the lower respiratory tract, ensuring consistent dosing and potentially enhancing immunogenicity at the primary site of pathogen entry. As such, the i.t. method represents an equally valuable, and in some respects advantageous, alternative to i.n delivery for evaluating mucosal vaccines in preclinical studies. Indeed, the i.t route of immunization serves as a relevant preclinical surrogate for aerosolized delivery methods currently being evaluated in human clinical trials for other infectious diseases, such as tuberculosis or COVID. Relatedly Jeyanathan et al (51) showed in a recent phase I clinical trial that individuals which had received a mRNA COVID-19 vaccine (Pfizer and/or Moderna) had, at baseline, no mucosal lung immunity (humoral or cellular), at least 3 months past their last dose. By contrast, the use of a Chimpanzee adenovirus platform (Ch-triCoV/Mac) for three SARS-CoV2 antigens was, compared to a human Ad-5 platform, efficient in filling the gap in respiratory immunity and generated, up to 48 weeks, measurable CD8 T cell (both circulating and resident memory TRM) and protective antibodies.

Overall, the present study demonstrates that our vaccine formulation, in addition to providing an advantageous auxotrophic phenotype adapted to the CF setting, was efficient, when given i.t, in preferentially inducing secretory IgA and TRMs at mucosal surfaces, a critical barrier that neutralizes pathogens before tissue invasion. We believe that it further underscores the clinical relevance of mucosal vaccination, and collectively support the utility of the i.t mouse model as a preclinical tool to investigate mucosal immunization strategies applicable to multiple delivery platforms.

## Acknowledgements

We wish to thank Mr Hugo Leroy and Mses Clémentine Cornu and Christelle Muñoz (our laboratory) for their help with some of the experiments and Mrs Geneviève Ball (Laboratoire de Chimie Bactérienne, UMR7283, Centre National de la Recherche Scientifique, Marseille) for generating the PAO1-Δ*lasB* mutant. We also thank Drs Loredana Saveanu (CRI/INSERM U1149, Paris) and Imane El Meouche (INSERM, IAME, Paris) for useful discussions. We are also indebted to Prs Catherine Llanes and Patrick Plésiat (Université Marie et Louis Pasteur, CNRS, Chrono-environnement (UMR 6249), Besançon, France)) for their gift of the *P.a* clinical strains (5-6 ; 8-20; 7-23; 9-8) and to *Vaincre la Mucoviscidose* and the association *Gregory Lemarchal* for continuous support. The authors declare that they have filed a patent application related to the technology described in this manuscript.

## Supplemental Figures legends

**Fig S1.**
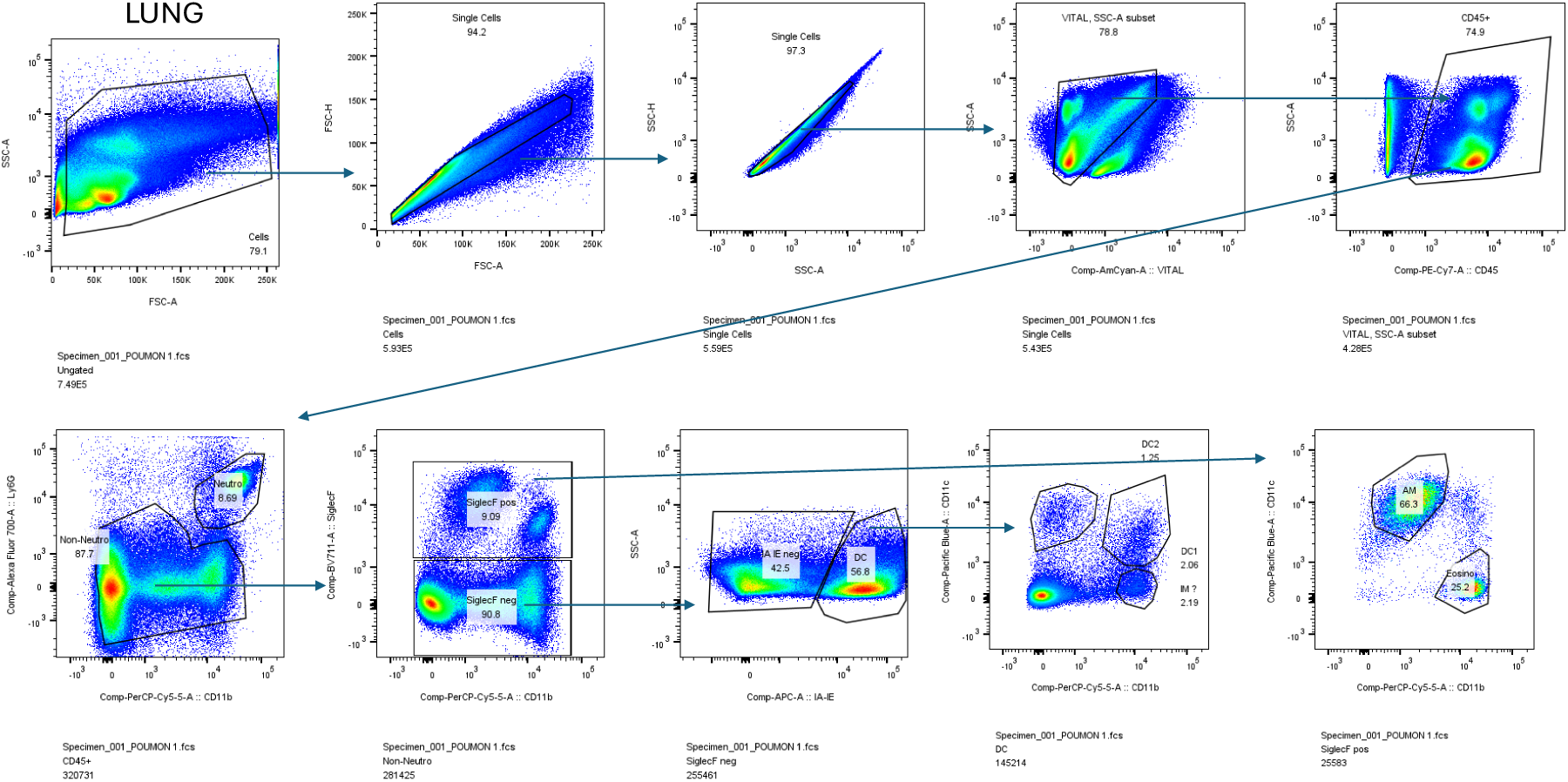
Gating strategy for the detection of lung myeloid cells

**Fig S2.**
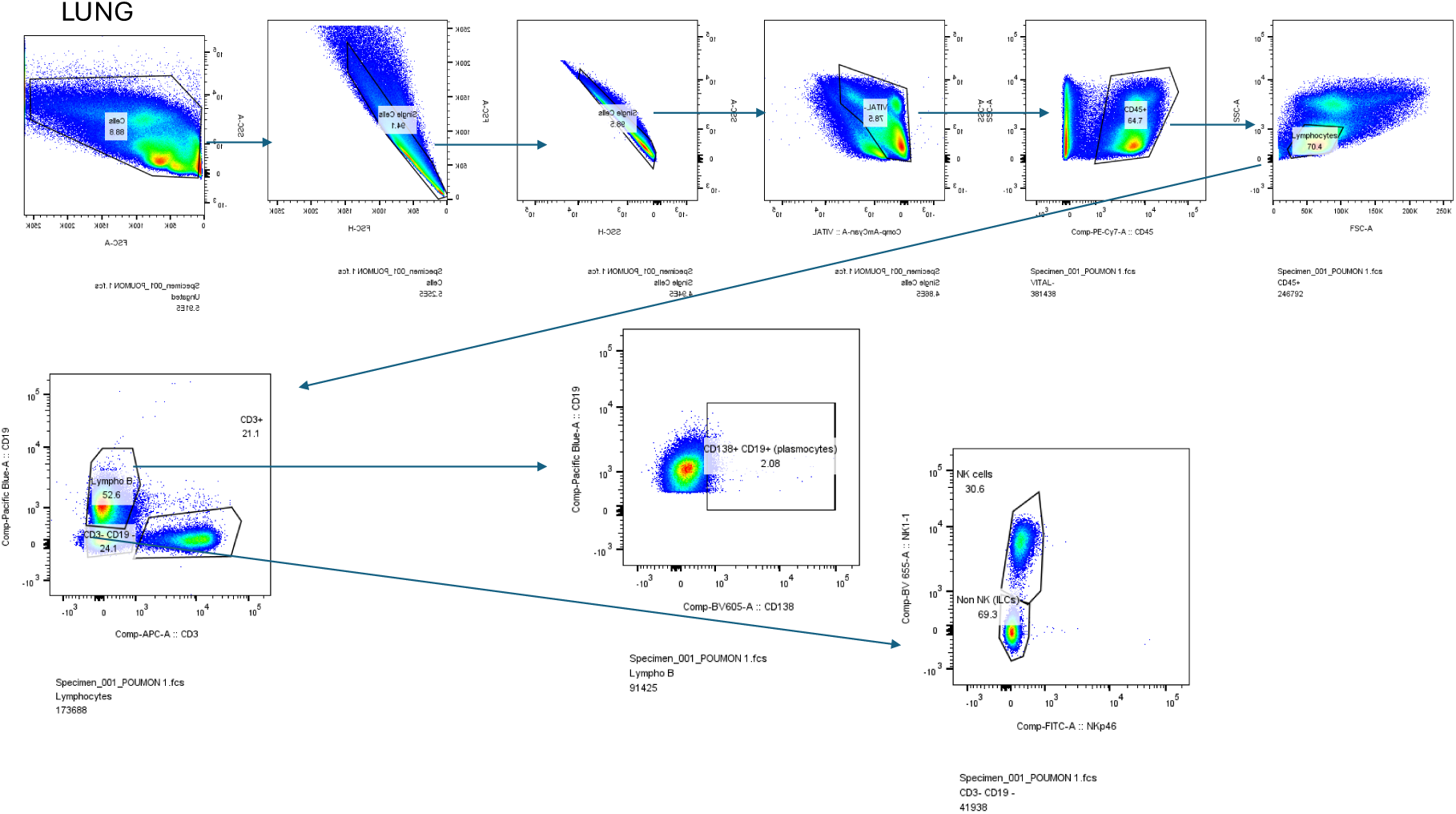
Gating strategy for the detection of lung lymphoid cells

**Fig S3.**
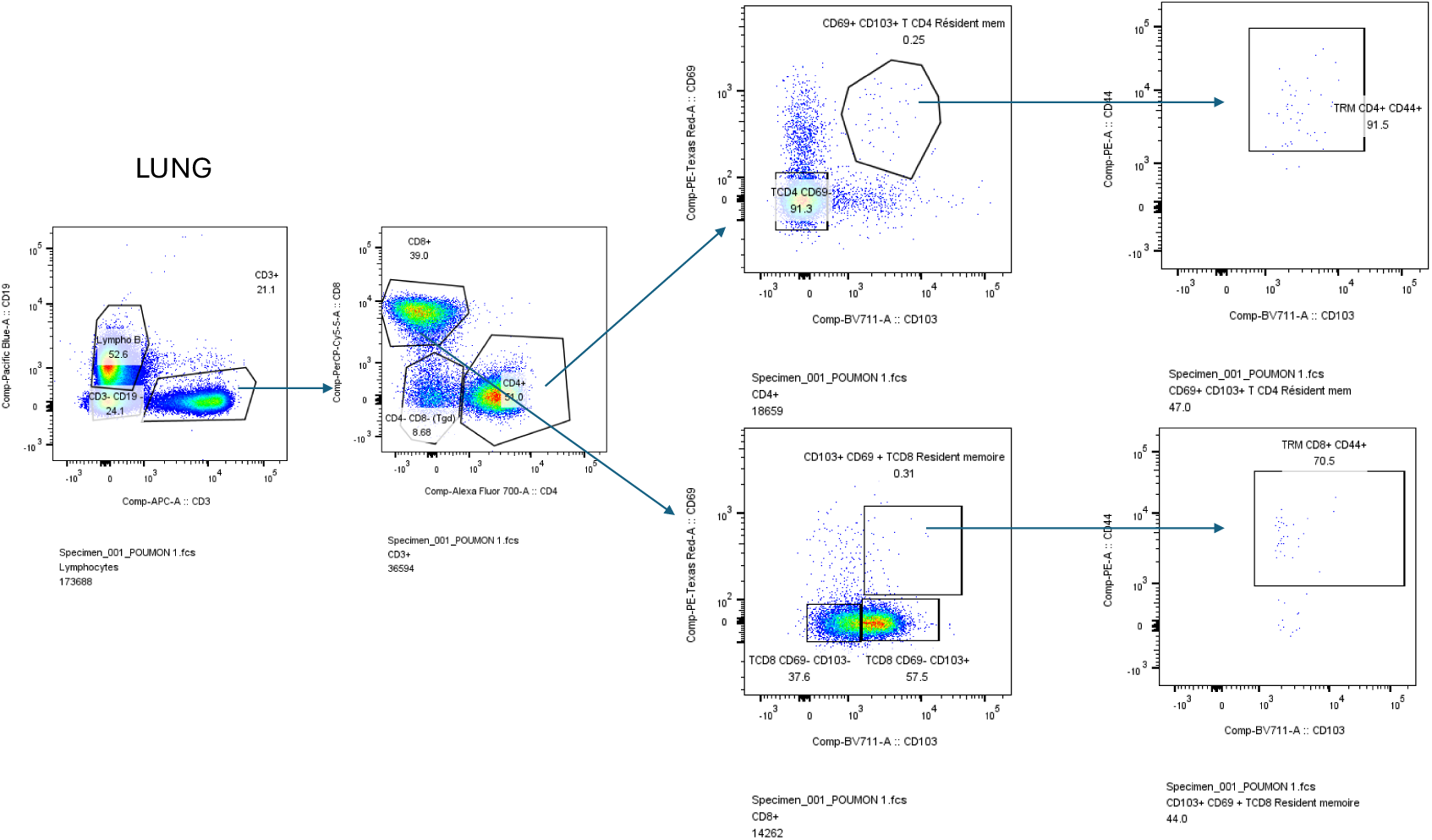
Gating strategy for the detection of lung CD4 and CD8 TRM (tissue resident memory cells)

**Fig S4:**
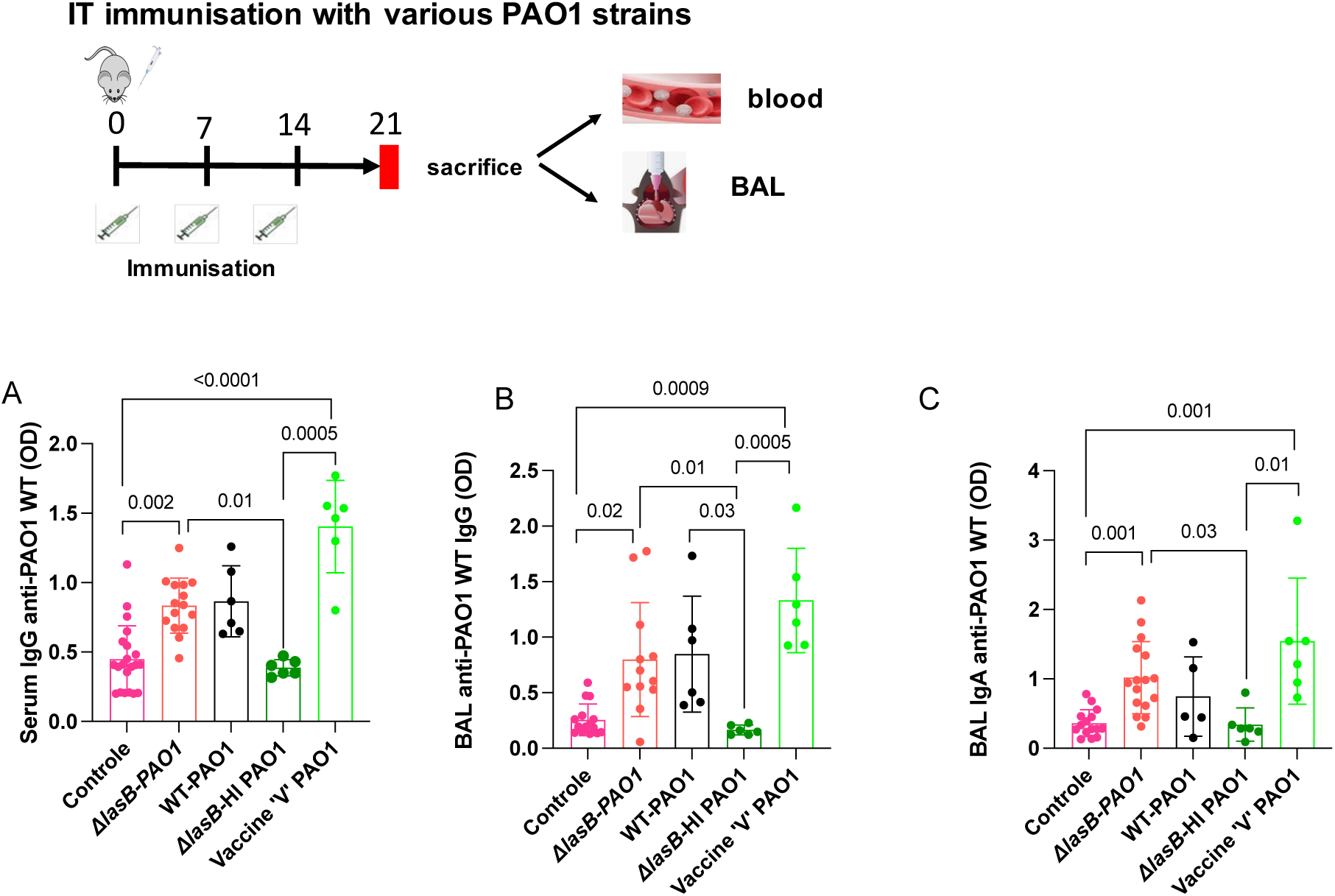
C57BL/6 male mice were given IT (at days 0, 7, 14) either NaCl (control) or were immunized (same time points) with either 10^6^ CFU of live ΔlasB-PAO1, live PAO1-WT, heat-inactivated (HI) ΔlasB-PAO1 or with the vaccine ‘V’ PAO1 strain. At day 21, 7 days after the last boost, blood and BALs were collected and used for anti-PAO1 serum total IgG (**A)**, anti-PAO1 BAL total IgG (**B)**, anti-PAO1 BAL IgA **(C).** Kruskal-Wallis Multiple comparaison test was used for to determine significance p<0.05. Error bars represent standard deviations (SD).

**Fig S5:**
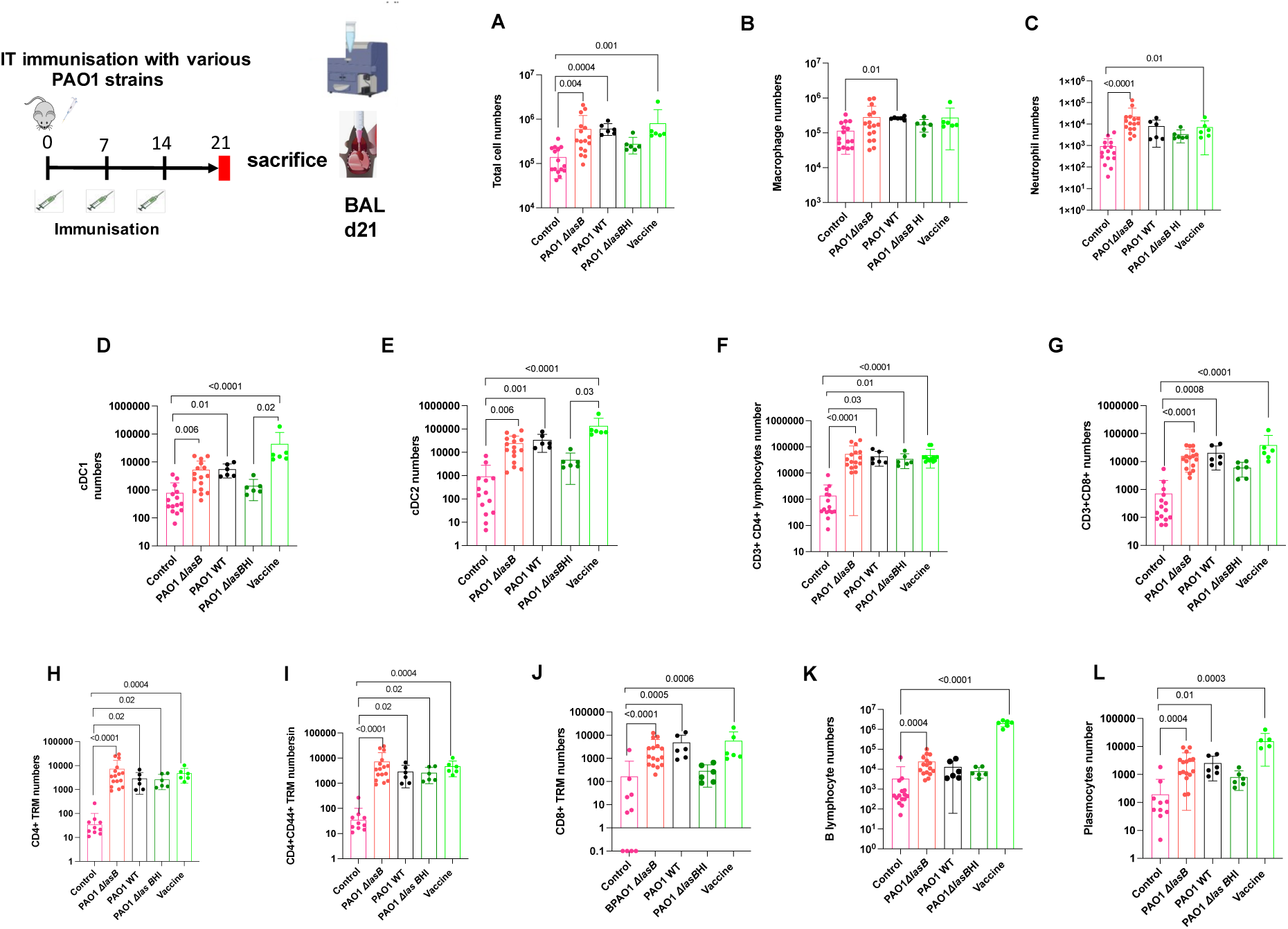
C57BL/6 male mice were immunized as in Fig S4 and BAL cells were recovered at d21 and analysed by FACS. Total cells, myeloid and lymphoid cells were quantified and levels plotted in panels A-L. Kruskal-Wallis Multiple comparaison test was used for to determine significance p<0.05. Error bars represent standard deviations (SD).

**Fig S6:**
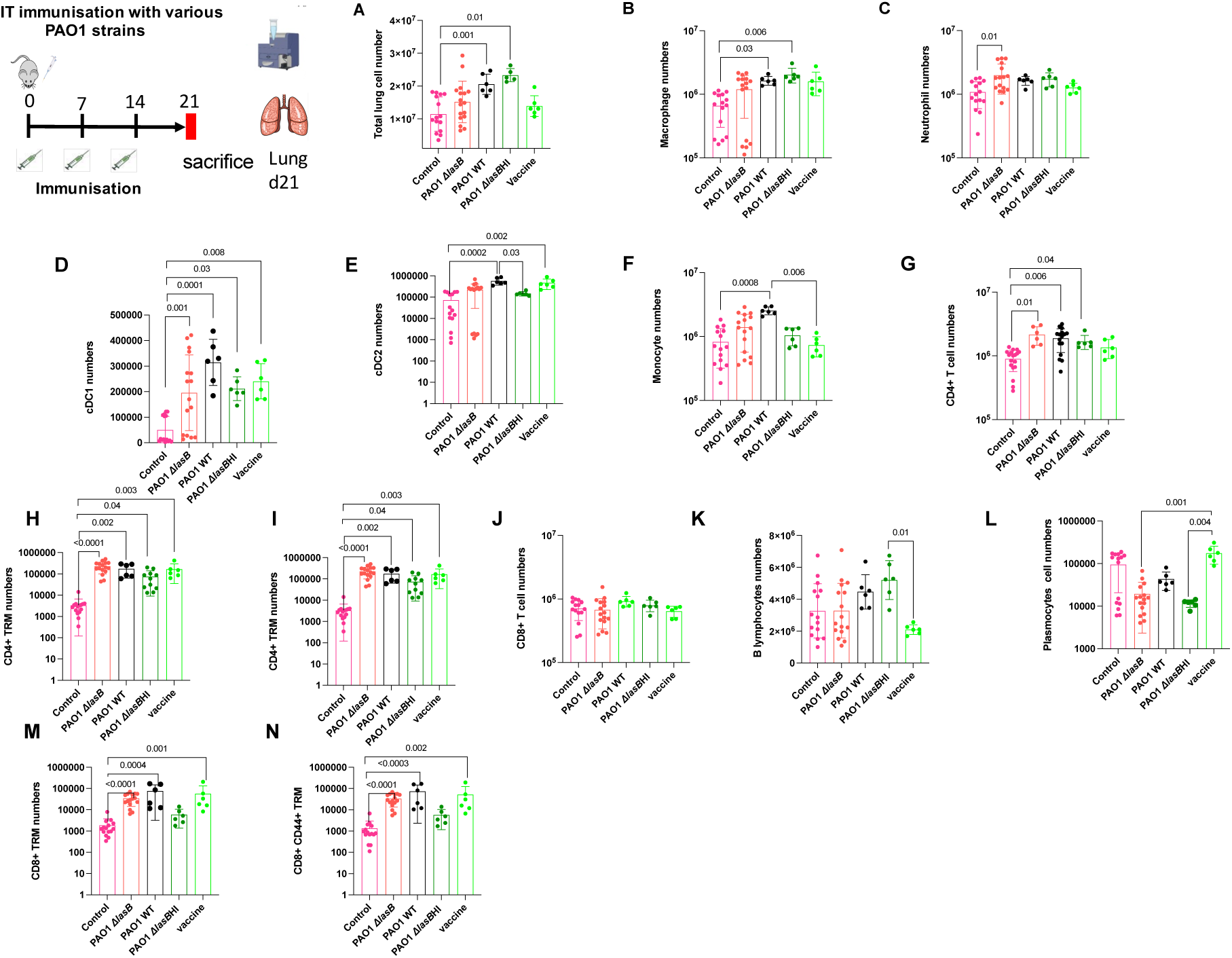
C57BL/6 male mice were immunized as in Fig S4 and lungs were recovered at d21. After homogenization, cells were analysed by FACS. Total cells, myeloid and lymphoid cells were quantified and levels plotted in panels A-L. Kruskal-Wallis Multiple comparaison test was used for to determine significance p<0.05. Error bars represent standard deviations (SD).

