## Supplementary material for "Mucosal vaccine immunity induced by a new auxotrophic *Pseudomonas aeruginosa* strain is linked to Th17 and IgA responses": Figs S1-S6

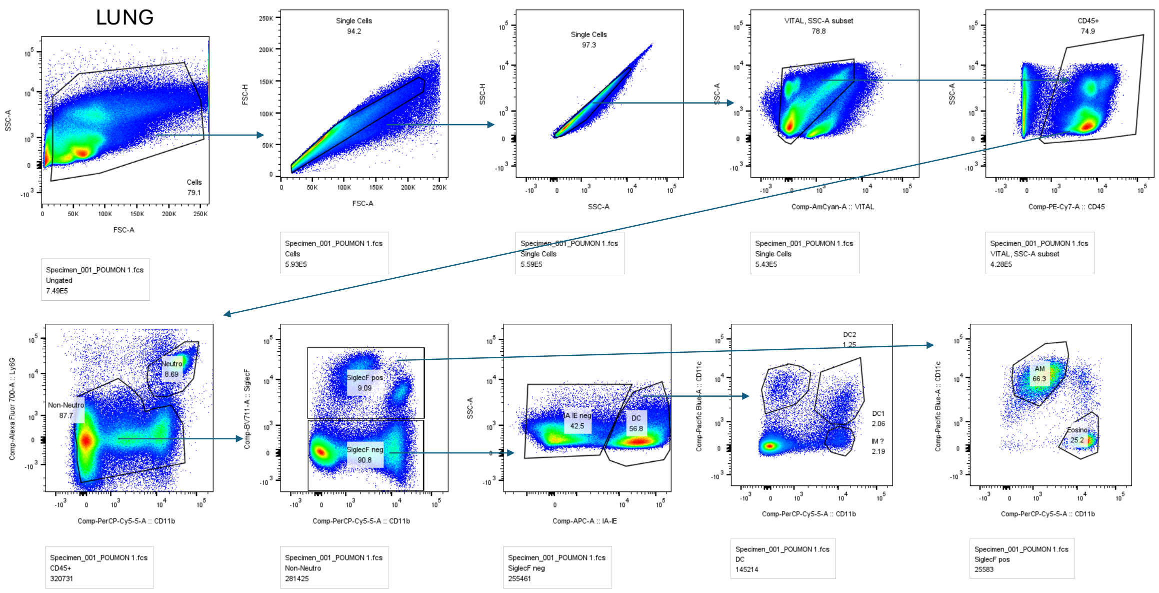
LUNG
Cells
79.1
Single Cells
94.2
Single Cells
97.3
VITAL, SSC-A subset
78.8
CD45+
74.9
FSC-A
SSC-A
FSC-H
SSC-H
Comp-AmCyan-A :: VITAL
Comp-PE-Cy7-A :: CD45
Specimen_001_POUMON 1.fcs
Ungated
7.49E5
Specimen_001_POUMON 1.fcs
Cells
5.93E5
Specimen_001_POUMON 1.fcs
Single Cells
5.59E5
Specimen_001_POUMON 1.fcs
Single Cells
5.43E5
Specimen_001_POUMON 1.fcs
VITAL, SSC-A subset
4.28E5
Neutro
8.69
Non-Neutro
87.7
SiglecF pos
9.09
SiglecF neg
90.8
IA IE neg
42.5
DC
56.8
DC2
1.25
DC1
2.06
IM ?
2.19
AM
66.3
Eosino
25.2
Comp-Alexa Fluor 700-A :: Ly6G
Comp-PerCP-Cy5-5-A :: CD11b
Comp-BV711-A :: SiglecF
Comp-APC-A :: IA-IE
Comp-Pacific Blue-A :: CD11c
Specimen_001_POUMON 1.fcs
CD45+
320731
Specimen_001_POUMON 1.fcs
Non-Neutro
281425
Specimen_001_POUMON 1.fcs
SiglecF neg
255461
Specimen_001_POUMON 1.fcs
DC
145214
Specimen_001_POUMON 1.fcs
SiglecF pos
25583

Fig S1

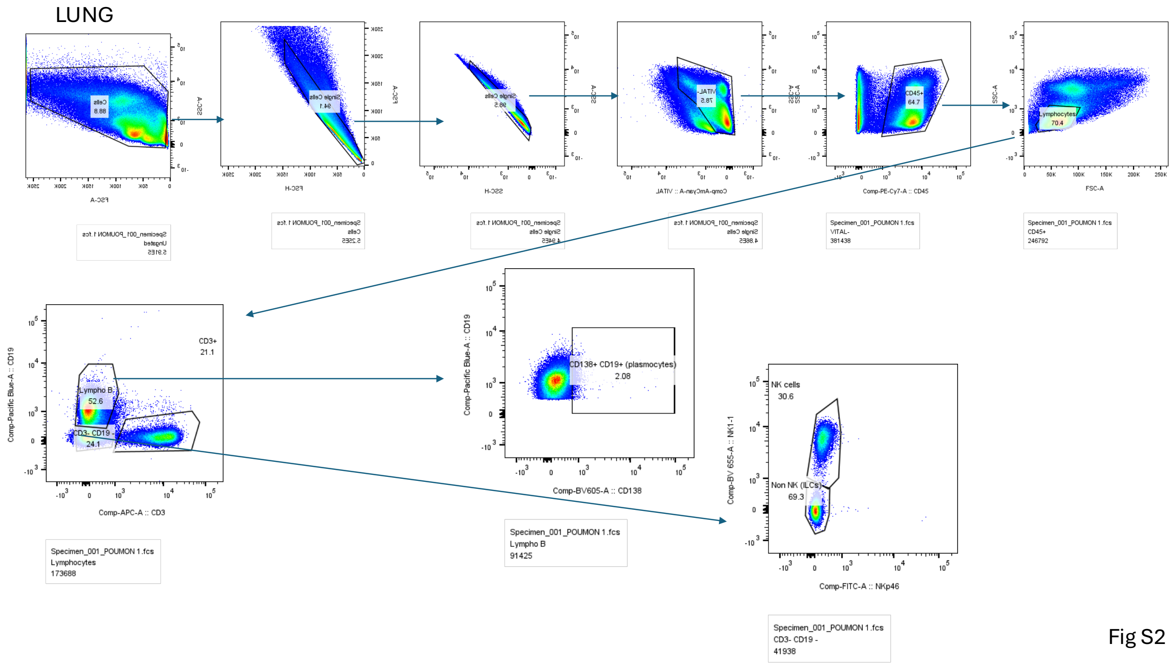
LUNG
CD45+
64.7
Lymphocytes
70.4
Comp-PE-Cy7-A :: CD45
FSC-A
SSC-A
Specimen_001_POUMON 1.fcs
VITAL-
381438
Specimen_001_POUMON 1.fcs
CD45+
246792
CD3+
21.1
Lympho B
52.6
CD3- CD19 -
24.1
Comp-Pacific Blue-A :: CD19
Comp-APC-A :: CD3
Specimen_001_POUMON 1.fcs
Lymphocytes
173688
CD138+ CD19+ (plasmocytes)
2.08
Comp-BV605-A :: CD138
Specimen_001_POUMON 1.fcs
Lympho B
91425
NK cells
30.6
Non NK (ILCs)
69.3
Comp-BV 655-A :: NK1-1
Comp-FITC-A :: NKp46
Specimen_001_POUMON 1.fcs
CD3- CD19 -
41938

Fig S2

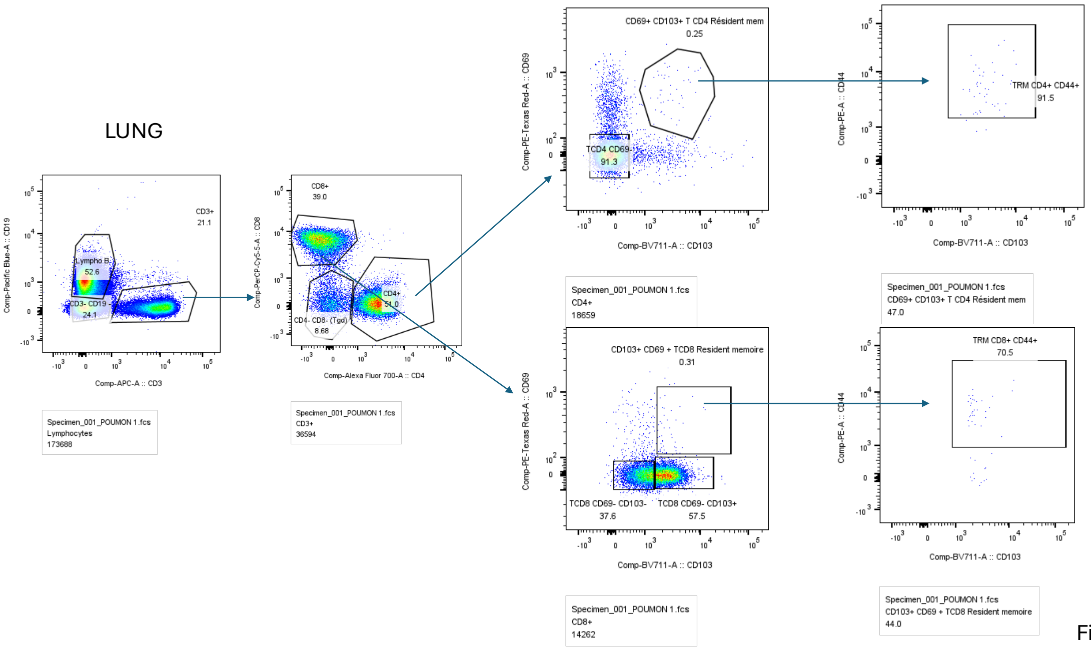
LUNG
Comp-APC-A :: CD3
Comp-Pacific Blue-A :: CD19
Lympho B
52.6
CD3+
21.1
CD3- CD19-
24.1
Specimen_001_POUMON 1.fcs
Lymphocytes
173688
Comp-Alexa Fluor 700-A :: CD4
Comp-PerCP-Cy5-5-A :: CD8
CD8+
39.0
CD4+
51.0
CD4- CD8- (Tgd)
8.68
Specimen_001_POUMON 1.fcs
CD3+
36594
CD69+ CD103+ T CD4 Résident mem
0.25
TCD4 CD69-
91.3
Comp-PE-Texas Red-A :: CD69
Comp-BV711-A :: CD103
Specimen_001_POUMON 1.fcs
CD4+
18659
TRM CD4+ CD44+
91.5
Comp-PE-A :: CD44
Specimen_001_POUMON 1.fcs
CD69+ CD103+ T CD4 Résident mem
47.0
CD103+ CD69 + TCD8 Resident memoire
0.31
TCD8 CD69- CD103-
37.6
TCD8 CD69- CD103+
57.5
Specimen_001_POUMON 1.fcs
CD8+
14262
TRM CD8+ CD44+
70.5
Specimen_001_POUMON 1.fcs
CD103+ CD69 + TCD8 Resident memoire
44.0

Fig S3

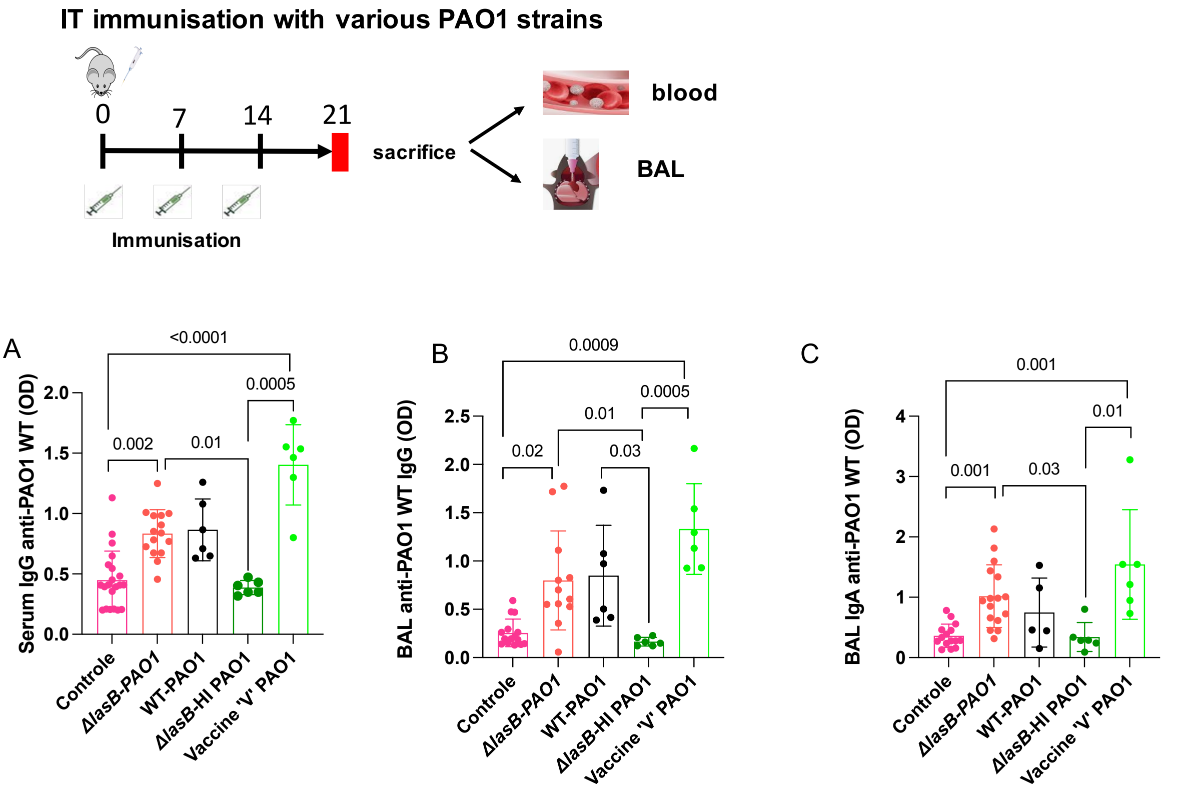
IT immunisation with various PAO1 strains
0
7
14
21
sacrifice
blood
BAL
Immunisation
A
Serum IgG anti-PAO1 WT (OD)
<0.0001
0.002
0.01
0.0005
B
BAL anti-PAO1 WT IgG (OD)
0.0009
0.02
0.01
0.03
0.0005
C
BAL IgA anti-PAO1 WT (OD)
0.001
0.001
0.03
0.01
Controle
ΔlasB-PAO1
WT-PAO1
ΔlasB-HI PAO1
Vaccine 'V' PAO1

Fig S4

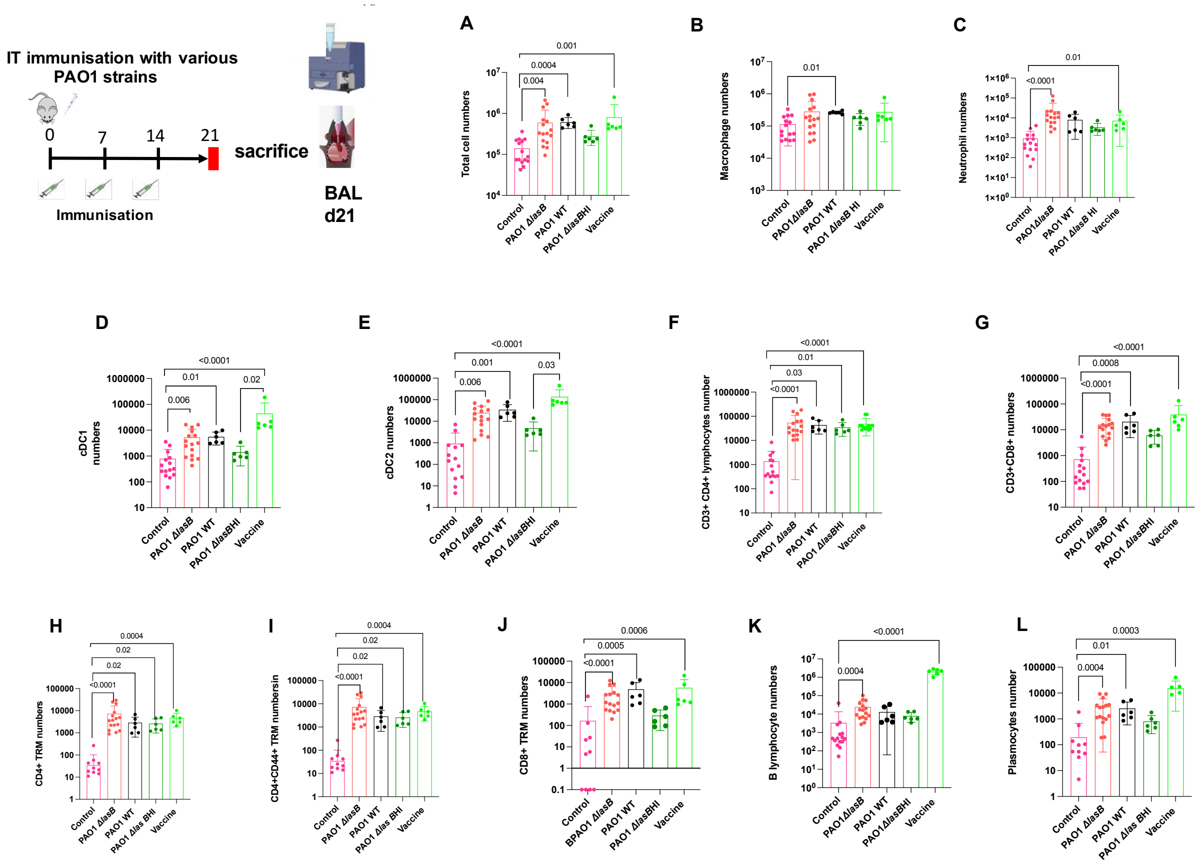
IT immunisation with various PAO1 strains
0
7
14
21
sacrifice
Immunisation
BAL d21
A
B
C
D
E
F
G
H
I
J
K
L
Total cell numbers
Macrophage numbers
Neutrophil numbers
cDC1 numbers
cDC2 numbers
CD3+ CD4+ lymphocytes number
CD3+CD8+ numbers
CD4+ TRM numbers
CD4+CD44+ TRM numbersin
CD8+ TRM numbers
B lymphocyte numbers
Plasmocytes number
Control
PAO1 ΔlasB
PAO1 WT
PAO1 ΔlasBHI
Vaccine

Fig S5

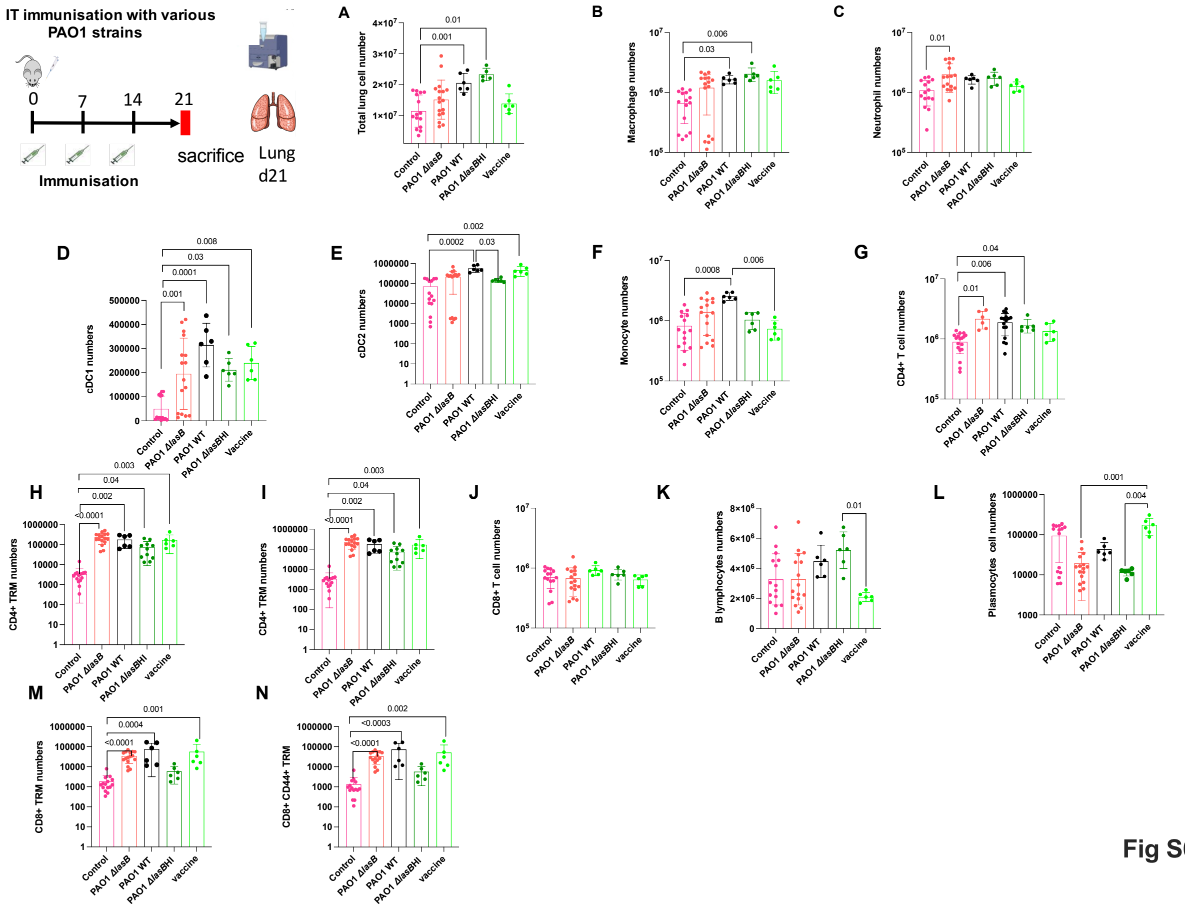

Fig S6

### Supplemental Figures legends

Fig S1Gating strategy for the detection of lung myeloid cells

Fig S2 Gating strategy for the detection of lung lymphoid cells

Fig S3 Gating strategy for the detection of lung CD4 and CD8 TRM (tissue resident memory cells)

Fig S4 : C57BL/6 male mice were given IT (at days 0, 7, 14) either NaCl (control) or were immunized (same time points) with either $10^6$ CFU of live ΔlasB-PAO1, live PAO1-WT, heat-inactivated (HI) ΔlasB-PAO1 or with the vaccine 'V' PAO1 strain. At day 21, 7 days after the last boost, blood and BALs were collected and used for anti-PAO1 serum total IgG (**A)**, anti-

PAO1 BAL total IgG (**B)**, anti-PAO1 BAL IgA **(C).** Kruskal-Wallis Multiple comparaison test was used for to determine significance $p<0.05$. Error bars represent standard deviations (SD).

Fig S5 : C57BL/6 male mice were immunized as in Fig S4 and BAL cells were recovered at d21 and analysed by FACS. Total cells, myeloid and lymphoid cells were quantified and levels plotted in panels A-L. Kruskal-Wallis Multiple comparaison test was used for to determine significance $p<0.05$. Error bars represent standard deviations (SD).

Fig S6 : C57BL/6 male mice were immunized as in Fig S4 and lungs were recovered at d21. After homogenization, cells were analysed by FACS. Total cells, myeloid and lymphoid cells were quantified and levels plotted in panels A-L. Kruskal-Wallis Multiple comparaison test was used for to determine significance $p<0.05$. Error bars represent standard deviations (SD).
